# Ventral Morphology and Ecological Implications of *Cindarella eucalla* (Artiopoda, Xandarellida) from Chengjiang Biota, China

**DOI:** 10.1101/2024.07.10.602865

**Authors:** Maoyin Zhang, Yu Liu, Huijuan Mai, Michel Schmidt, Xianguang Hou

## Abstract

Artiopoda, an early euarthropod group, displays post-antennal appendages resembling trilobite limbs, but relationships with other early euarthropods remain enigmatic. Limited studies and morphological details also hinder the understanding of internal relationships within Artiopoda. Recently, exceptionally well preserved euarthropod fossils from the Chengjiang Biota were studied using X-ray computed tomography, revealing detailed morphologies. In this study, *Cindarella eucalla*, a xandarellid from the Chengjiang Biota, was re-investigated using X-ray computed tomography and fluorescent microscopy. We document a posterior extension of the head shield over the anterior six trunk tergites, axial spines on the posterior trunk, a natant hypostome, four pairs of post-antennal cephalic appendages, and a dorsoventral mismatch between posterior tergites and limb pairs. Under parsimony, Xandarellida is recovered as monophyletic; Bayesian analysis also recovers the clade, but with moderate basal support (PP = 0.55). Similarities between the appendages of *C. eucalla* and those of selected fossil and extant arthropods are described here as morphological comparisons only. These anatomical data provide a basis for future tests of xandarellid phylogeny and palaeoecology.

## Introduction

Artiopoda is an extinct group of early euarthropods containing Vicissicaudata, Protosutura, Trilobita, and other non-biomineralized euarthropods (mainly referring to Conciliterga, Nectopleura, Petalopleura and other euarthropods with similar dorsal exoskeleton morphology to that of trilobites) (Hou and Bergström, 1997; Du et al., 2019; Mayers et al., 2019). Some analyses have suggested that Artiopoda is paraphyletic with respect to Panchelicerata and total-group Mandibulata (Aria, 2022), whereas others resolved Artiopoda as the sister group to Panchelicerata within total-group Chelicerata (Ortega-Hernández et al., 2013), and still other studies regarded it as the sister group to total-group Mandibulata (Stein et al., 2013; Aria et al., 2020; Zeng et al., 2020). Opinions on the relationship of animals within and out of this clade vary (Lerosey-Aubril et al., 2017; Schmidt et al., 2022; Zhang et al., 2022a; Zhu et al., 2023). While some artiopodans are known from complete dorsal exoskeletons and even articulated appendages, such exceptional preservation is largely restricted to Konservat-Lagerstätten. With the exception of heavily biomineralized trilobites, most non-biomineralized artiopodans are typically preserved only as incomplete dorsal exoskeletons (Ortega-Hernández et al., 2013; Hou et al., 2017; Berks et al., 2023). In recent years, detailed ventral morphology of several artiopodans, such as *Sinoburius lunaris* (Chen et al., 2019; Schmidt et al., 2021), *Naraoia spinosa* (Zhai et al., 2019), *Retifacies abnormalis* (Zhang et al., 2022a), *Pygmaclypeatus daziensis* (Schmidt et al., 2022, 2025), and *Sidneyia minor* (Du et al., 2023), were revealed using X-ray computed tomography. *C. eucalla* has previously been interpreted as a possible predator or scavenger, partly on the basis of its conspicuous compound eyes (Hou and Bergström, 1997; Ramsköld et al., 1997; Hou et al., 2017; Chen et al., 2019). Such functional interpretations remain hypotheses because eye size alone does not establish visual performance or feeding mode. The ventral anatomy and phylogenetic relationships of *C. eucalla* are also in-completely known (Ramsköld et al., 1997; Legg et al., 2013; Hou et al., 2017; Chen et al., 2019; Du et al., 2019; Berks et al., 2023; Zhu et al., 2023). Here, we use X-ray computed tomography and three-dimensional visualisation to document previously inaccessible morphology and to evaluate its phylogenetic placement.

## Results

### Systematic Palaeontology

Euarthropoda Lankester, 1904

Artiopoda Hou & Bergström, 1997

Xandarellida Chen, Ramsköld, Edgecombe & Zhou in Chen et al., 1996

#### Constituent taxa

*C. eucalla* Chen, Ramsköld, Edgecombe & Zhou in Chen et al., 1996 (Chen et al., 1996; Chen, 2004; Hou et al., 2017); *Xandarella spectaculum* Hou, Ramsköld & Bergström, 1991 (Hou et al., 1991; Hou and Bergström, 1997; Hou et al., 2017); *Sinoburius lunaris* Hou, Ramsköld & Bergström, 1991 (Hou et al., 1991; Chen et al., 2019; Schmidt et al., 2021); *Luohuilinella rarus* Zhang, Fu, & Dai, 2012 (Zhang et al., 2012); *Luohuilinella deletres* Hou et al., 2018 (Hou et al., 2018); *Zhugeia acuticaudata* Zhu et al., 2023 (Zhu et al., 2023).

#### Emended diagnosis

Examination of the available materials reveals that this taxon is mostly preserved in a dorsoventrally flattened state, with a broad, semicircular head shield that has stalked eyes originating from the ventral side. The head shield has dorsal exoskeleton bulges or notches that accommodate the eyes, or the eyes simply extend out from the lateral edge of the head shield. The natant hypostome, a pair of uniramous antennae, and three to five pairs of biramous appendages are covered by the head shield. In some genera, the antenna bears one to two antennal scales. The head shield extends posteriorly, either covering several reduced thoracic tergites or followed immediately by a reduced one. The middle and/or posterior trunk tergites have variable patterns of dorsoventral mismatch relative to the ventral biramous appendage pairs. Each trunk tergite covers more than one limb pair. Endopodal endites are spiny, small, or entirely absent (Ramsköld et al., 1997; Hou et al., 2017, 2018; Chen et al., 2019).

#### Remarks

Five genera comprise this taxon: *Cindarella* (*Chen et al., 1996*; *Ramsköld et al., 1997*; *Edgecombe and Ramsköld, 1999*; *Hou et al., 2017*), *Xandarella* (*Hou and Bergström, 1997*; *Ramsköld et al., 1997*; *Edgecombe and Ramsköld, 1999*; *Hou et al., 2004*, *2017*), *Luohuilinella* (*Zhang et al., 2012*; *Hou et al., 2018*), *Sinoburius* (*Chen et al., 2019*; *Schmidt et al., 2021*), *Zhugeia* (*Zhu et al., 2023*). *Almenia spinosa* Hou and Bergström, 1997 was previously considered to be synonymous with *C. eucalla* (*Hou and Bergström, 1997*; *Ramsköld et al., 1997*; *Edgecombe and Ramsköld, 1999*; *Hou et al., 2004*, *2017*). However, our analysis suggests that *A. spinosa* is actually synonymous with *X. spectaculum*, not *C. eucalla*. Here follows the reasons:

1. The edges of *A. spinosa*’s head shield are relatively sharp, and it has posterolateral corner that extends into a pointed genal angle — a feature more similar to *X. spectaculum* than *C. eucalla*, which has a blunt and rounded head shield without a distinct genal angle.
2. The number of post-antennal cephalic appendage pairs in *A. spinosa*—six or more—is consistent with *X. spectaculum*, unlike *C. eucalla*, which has only four pairs.
3. *A. spinosa* has eleven trunk tergites, possibly followed by a small tergite close to the head shield—almost the same to *X. spectaculum*. In contrast, *C. eucalla*’s head shield covers six anterior trunk tergites, followed by 16 to 26 trunk tergites.
4. The last four tergites of *A. spinosa* are subequal in length, gradually lengthening towards the rear (as seen in Hou and Bergström, 1997, Figures 75, 76). This pattern is similar to that of *X. spectaculum*, whereas all trunk tergites of *C. eucalla* are nearly equal in length.

3D anatomical data reveals that *Xandarella mauretanica* (Ortega-Hernández et al., 2017a) is actually a trilobite *Gigoutella mauretanica* (El Albani et al., 2024), rendering *Xandarella mauretanic* an invalid species. Currently, only the dorsal exoskeleton of *Austroxandarella poikar* has been reported (Edgecombe et al., 2025). Although the progressive increase in width across the last four tergites likely represents a characteristic of Xandarellida, the dorsal exoskeleton shows no evidence of eyes, and the head shield lacks both eye slits and eye notches; furthermore, no small tergite has been identified. Accordingly, we do not assign this species to Xandarellida at present, leaving this question open to future reassessment should more conclusive evidence become available.

The dorsal exoskeleton bulges or eye notches on head shield have been reported in *X. spectaculum* (Hou et al., 1991; Hou and Bergström, 1997; Hou et al., 2017), *Sinoburius lunaris* (Hou et al., 1991; Hou and Bergström, 1997; Hou et al., 2017; Chen et al., 2019; Schmidt et al., 2021), *Luohuilinella ramus* (Zhang et al., 2012), and *L. deletres* (Hou et al., 2018), but not in *C. eucalla* (Chen et al., 1996; Chen, 2004; Hou et al., 2017). Natant hypostome have been reported in all xandarellid species except *L. ramus* (Hou et al., 1991; Hou and Bergström, 1997; Luo et al., 1997; Hou et al., 1999, 2004, 2017, 2018).

In the systematic paleontology of Chen et al. (2019), *Phytophilaspis pergamen*a was classified within xandarellida, and the phylogenetic analysis shows that this species clusters with other taxa in the xandarellida lineage (Chen et al., 2019, Fig. 9). Morphological comparisons reveal the following key distinctions:

1. The facial sutures of *P. pergamena* (Ivantsov, 1999) superficially resemble those of xandarellids (such as *X. spectaculum* (Hou et al., 1991, 2017), *L. ramus* (Zhang et al., 2012), *L. deletres* (Hou et al., 2018), *S. lunaris* (Hou et al., 1991, 2017; Hou and Bergström, 1997). In reality, *P. pergamena* has no eye notch, and the facial sutures do not intersect or border the ocular area. This morphology is fundamentally different from the cephalic structures associated with visual area in related taxa: *L. deletres* (Hou et al., 2018), *L. ramus* (Zhang et al., 2012) and *S. lunaris* (Chen et al., 2019; Schmidt et al., 2021) possess an eye notch (an inward excavation of the head shield’s lateral margin forming a recess for eye accommodation, or an opening directly on the dorsal surface that houses the eye), whereas *X. spectaculum* (Hou et al., 1991; Hou and Bergström, 1997; Hou et al., 2017) exhibits a more derived eye slit. This slit forms when the inward excavation of the notch is so pronounced that its anterior and posterior margins overlap.
2. Both *P. pergamena* and xandarellids possess dorsally positioned eyes, this appears to be a superficial similarity. In xandarellids, the eyes are ventral stalked eyes that project dorsally through the head shield. For *P. pergamena*, the nature of its dorsal eyes remains uncertain; there is currently no conclusive evidence to determine whether they are stalked eyes extending from the ventral side or sessile eyes fixed directly on the dorsal surface.
3. The thoracic tergites of *P. pergamena* lack distinct pleurae. Its pygidium is bluntly rounded terminally, lacking clear lateral or terminal spines. This overall morphology more closely resembles that of the subclass Nektopleura or Conciliterga than that of Xandarellida.
4. Our phylogenetic analysis did not resolve *Phytophilaspis pergamena* within the xandarellid clade.

Based on the combination of these morphological and phylogenetic considerations, we have not assigned *P. pergamena* to the Order Xandarellida in this study.

No antennal scale was mentioned in previous studies or in this investigation on *X. spectaculum* (Hou et al., 1991; Hou and Bergström, 1997; Hou et al., 2017), *L. ramus* (Zhang et al., 2012) and *L. deletres* (Hou et al., 2018) but one on the inner side of *S. lunaris* (Chen et al., 2019; Schmidt et al., 2021) and *C. eucalla* revealed in this study. The number of post-antennal cephalic appendage pairs ranges from three to five: Three pairs in *L. deletres* (Hou et al., 2018), four pairs in *C. eucalla* and *S. lunaris* (Chen et al., 2019; Schmidt et al., 2021), five pairs in *X. spectaculum* (Hou et al., 1991; Hou and Bergström, 1997; Hou et al., 2017). The head shield in *X. spectaculum* (Hou et al., 1991; Hou and Bergström, 1997; Hou et al., 2017), *L. ramus* (Zhang et al., 2012), *L. deletres* (Hou et al., 2018) and *S. lunaris* (Chen et al., 2019; Schmidt et al., 2021) is closely followed by a small tergite without pleural elongations, but in *C. eucalla*, the head shield extended posteriorly like a folded carapace to cover the anterior six trunk tergites. The dorsoventral segmental mismatch between trunk tergites and ventral limb pairs displayed various patterns:

1. The last four tergites lengthened with an increase in appendage numbers posteriorly in *X. spectacunlum*;
2. The last thoracic tergite covers two ventral appendage pairs in *S. lunaris* (Chen et al., 2019; Schmidt et al., 2021);
3. Posterior to the sixth trunk tergite, *C. eucalla* displays a gradual increase in the number of appendage pairs underlying each tergite; the tergite margins (dorsal) no longer align with the appendage pair boundaries (ventral) (Hou et al., 2017).

Consequently, *Almenia spinosa*, *Xandarella mauretanica*, *Phytophilaspis pergamena* are definitively excluded from Xandarellida, while *Austroxandarella poikar* is provisionally omitted, pending further evidence of its ventral morphology and phylogenetic placement.

The endites on the inner side of endopods vary among taxa: *S. lunaris* lacks endites but possesses gnathobases with blunt spines on the proximal protopods (Chen et al., 2019; Schmidt et al., 2021); in *X. spectaculum*, endites on endopod podomeres are indistinct (Hou and Bergström, 1997; Hou et al., 2017); *C. eucalla* bears a cluster of blunt spines on each endopod podomere; and *L. deletres* has a pair of sharp spines on each endopod podomere (Hou et al., 2018). *S. lunaris* (Schmidt et al., 2021) and *X. spectaculum* (Hou et al., 2017) possess distinct pygidia that are larger than their preceding tergites, whereas *L. ramus* (Zhang et al., 2012), *L. deletres* (Hou et al., 2018) and *C. eucalla*, although theoretically possessing pygidia, lack a well-developed or enlarged pygidium and show no such morphological distinction. Individuals with distinctive large pygidia have a median spine and two lateral spine pairs (e.g., in *S. lunaris* and *X. spectaculum*), and four (in *C. eucalla*) or more (in *L. deletres*) axial spines on posterior tergites in these who lack an obvious pygidium. Therefore, we emend the diagnosis of Xandarellida as presented above.

### Genus Cindarella Chen, Ramsköld, Edgecombe & Zhou in Chen et al., 1996

#### Emended diagnosis

The body is divided into two tagmata, the cephalon and the trunk, with the anterior six tergites of the latter covered by extensions of the head shield. The dorsal exoskeleton of the trunk, consisting of 22 to 32 tergites, is spindle-shaped, with the ninth tergite being the widest. The terminal four tergites show extensions of median ridge into long wide spines. All tergites are equal in length. *C. eucalla* has a pair of large, stalked compound eyes. The cephalic appendages include a pair of multi-annulate antennae and four pairs of biramous appendages with a similar morphology to those of the trunk. The cephalic appendages become robust and larger from anterior to posterior. Each of the anterior six tergites in the trunk covers a pair of biramous appendages, and each of the following tergites covers more than one pair, indicating a decoupling of tergites and ventral appendage pairs. All trunk appendages are biramous and homonomous. Each biramous appendage consists of a protopod, a six-podomere endopod with a cluster of spines on each podomere, an exopod with dozens of lamellae on a paddle-like shaft, and a rod-like exite.

### Cindarella eucalla Chen, Ramsköld, Edgecombe & Zhou in Chen et al., 1996

1996 *Cindarella eucalla* Chen et al., Taiwan Museum of Natural Science, pp. 160–162, Figures 208–210.

1997 *Almenia spinosa* Hou and Bergström, Scandinavian University Press, pp. 81–83, Figures 71–73.

1997 *Cindarella eucalla* Ramsköld et al., Transactions of the Royal Society of Edinburgh-Earth Sciences, Figures 1–14.

**Figure 1.**
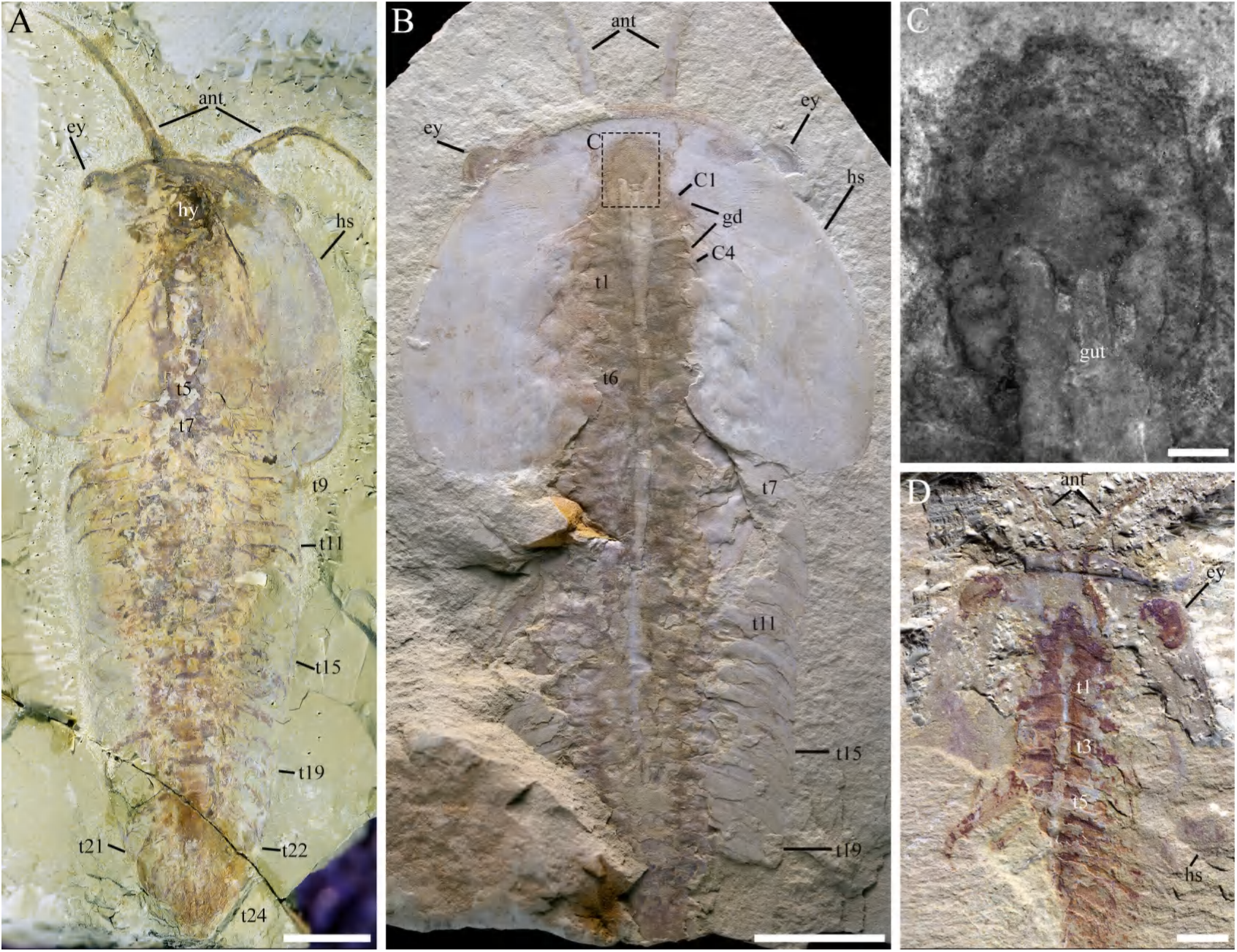
Macro-photographic (A, B, D) and fluorescent (C) images of *C. eucalla*. **A** Dorsal view of YKLP 17360. **B** Dorsal view of YKLP 17363a. **C** Close-up of hypostome in **B**. **D** Ventral view of YKLP 17373. Abbreviations: ant, antenna(e); C*n*, Post-antenna cephalic appendage number; el, exopod lobe (shaft); exo, exopod; ey, eye(s); gd, gut diverticula(e); gut, gut content/trace; hs, head shield; t*n*, tergite number. Scalebar: 1 cm in **A** and **B**, 2 mm in **D**, 1 mm in **C**.

1999 *Cindarella eucalla* Luo et al., Yunnan Science and Technology Press, pp. 49, plate 6, Figure 4

2004 *Cindarella eucalla* Hou et al., Blackwell Publishing Ltd., pp. 170–171, Figures 16.57–16.58. 2013 *Cindarella eucalla* Zhao et al., Scientific Reports, Figures 1–2.

**Figure 2.**
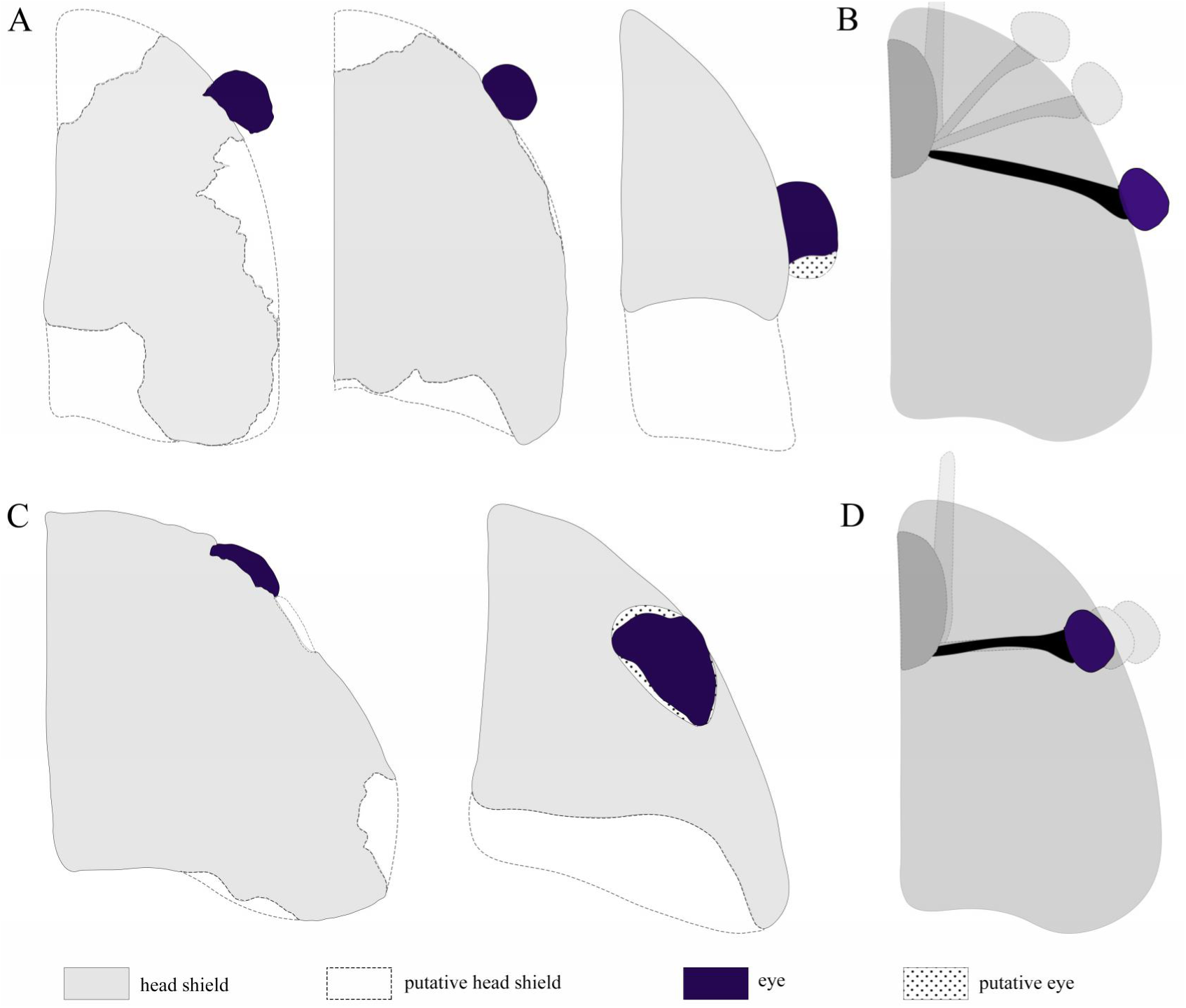
Eye-position variability in preserved specimens of *C. eucalla*. Gray area indicates head shield, dashed line indicates putative head shield, deep purple area indicates eye, doted area indicates putative eye in **A** and **C**. **A** Outline of head shield and eyes in three specimens of *C. eucalla* (Left to right): YKLP 17371, YKLP 17368 and YKLP 17366. **B** Schematic diagram of eye positions in **A**. **C** Outline of head shield and eyes in two specimens of *C. eucalla* (Left to right): YKLP 17365, YKLP 17368. **D** Schematic diagram of eye positions in **A** and **C**. **B** and **D** show alternative positional reconstructions only; they are not reconstructions of an eye-movement trajectory. Positional differences may reflect biological mobility, taphonomic displacement, or both. Not to scale.

2017 *Cindarella eucalla* Hou et al., John Wiley & Sons Ltd., pp. 206–207, Figures 20.37–20.38. Diagnosis: As for genus.

### Description

#### Dorsal exoskeleton

The exoskeleton is non-biomineralized and most specimens preserved dorsoventrally flattened (Figures 1 A, B, D; 2–5; 6 A; 7 A, C; 8 A; Figure 1—figure supplement 1; Figure 6—figure supplement 6 A, C), only a few specimens preserved laterally (YKLP 17369 in Fig. 3 and Figure 3—figure supplement 3, YKLP 17366 in Figure 2—figure supplement 2 A; supplementary video 1, 2). Complete specimens examined herein (appendages excluded) measure 11.60-–90.25 mm in sagittal length, from the anterior margin of the cephalon to the posterior end of the last tergite. However, specimens up to 130 mm have previously been reported in Hou et al. (2017). The dorsal exoskeleton is divided into two tagmata: a large semi-elliptical head shield and the trunk with 22 to 32 tergites (ELRC 18502 shows 22 tergites, ELRC 18505 shows 24 tergites in (Ramsköld et al., 1997); YKLP 17371 in Figure 7 D shows 23 tergites; YKLP 17360 in Figure 1 A shows 24 tergites; YKLP 17374 in Figure 6—figure supplement 6 C shows 28 tergites; YKLP 17361 in Figure 6 A shows 32 tergites). No ornaments is observed on the dorsal exoskeleton (Figures 1 A, B; 6 A, 7 A, C; Figure 1—figure supplement 1 A, B; Figure 2—figure supplement 2 A–F; Figure 6—figure supplement 6 C). The head shield has no sharp genal spines but blunt lateral margins (Figures 1 A, B; 3 A; 6 A; 7 A, C; 8 A; Figure 1—figure supplement 1 A, B; Figure 2—figure supplement 2 B; Figure 6—figure supplement 6 C). Its length (measured from the anterior margin to the posterior margin of head shield) is 33.68%–46.89% of the body length, ranging from 8.79 to 59.80 mm in this study. The maximum width (which is also the width of the head shield) is 84.17%–145.30% of the length, ranging from 10.46 to 71.86 mm. The posterior extension of the head shield coveres the first six trunk tergites (Figures 1 A, B, D; 3 A, C; 6 A, B; Figure 1—figure supplement 1 A, B; Figure 6—figure supplement 6 C). The dorsal trunk exoskeleton is spindle shaped, with the dorsal nine tergites tapering anteriorly and the others tapering posteriorly. All tergites are in equal lengths (Figures 1 A, C; 3 A; 6 A; 7 A, C; Figure 1—figure supplement 1 A, B; Figure 2—figure supplement 2 A, B, F; Figure 6—figure supplement 6 C). Median ridges of the last four tergites extended into axial spines (Figure 7 B, D).

**Figure 3.**
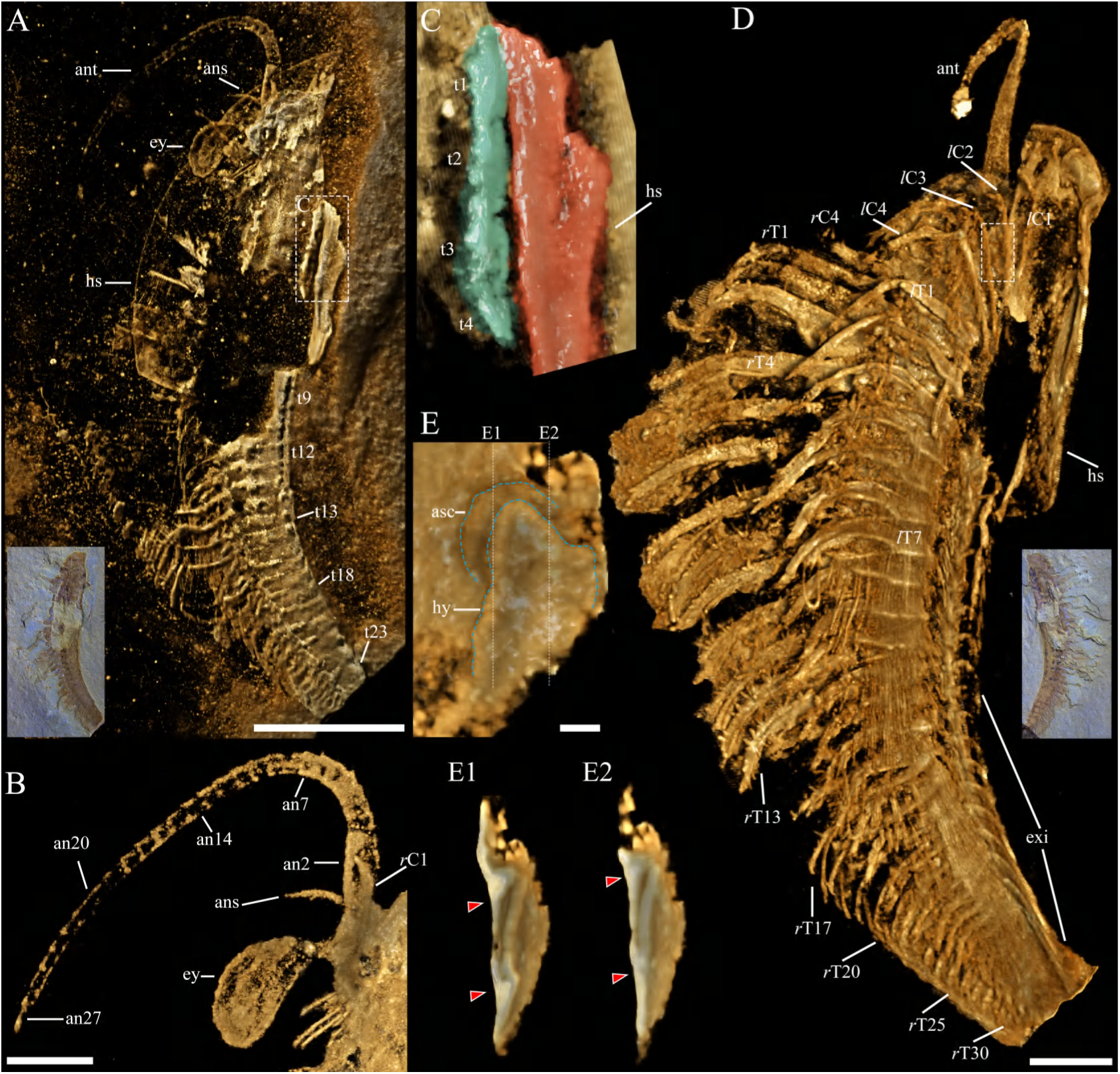
Digital anatomy of *C. eucalla* (YKLP 17369) based on X-ray tomographic data processed in Drishti v2.4. **A** YKLP 17369a, dorsal view. **B** YKLP 17369a, left antenna and eye, dorsal view. **C** overlap of head shield (in red) and trunk tergites (in cyan) within the dashed box in **A**. **D** YKLP 17369b, Ventral view, body and appendages digitally extracted from rocks. **E** Anterior sclerite and hypostome within the dashed box in **D** with the second and third left cephalic appendages romoved, ventral view (flipped horizontally). The dashed lines indicate two longitudinal sections in **E1** and **E2**. **E1**–**E2** two longitudinal sections (flipped horizontally) indicated in **E**. The arrowheads indicate separated sclerites posited anterior to first post-antennal appendage pair. Abbreviations: an*n*, antennal segment number; ant, antenna(e); ans, antennal scale; asc, anterior sclerite; C*n*, post-antennal cephalic appendage number; exi, exite(s); ey, eye(s); hs, head shiled; hy,hypostome; t*n*, tergite number; T*n*, trunk appendage number; *l*, left; *r*, right. Scalebar: 5 mm in **A**, 2 mm in **D**, 1 mm in **B**, 0.2 mm in **E**, no scalebar in **C**, **E1** and **E2**.

**Figure 4.**
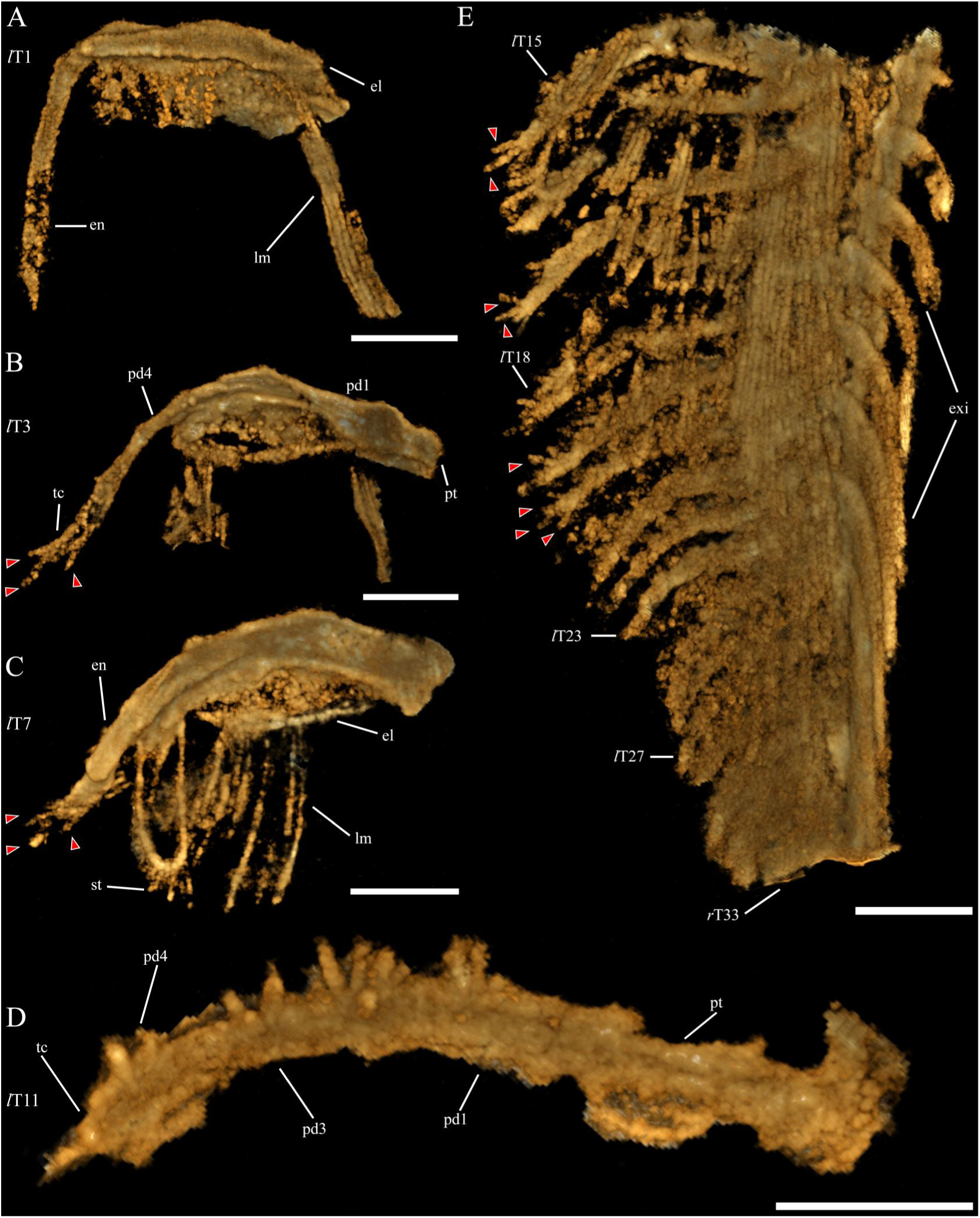
Digital anatomy of *C. eucalla* (YKLP 17369b) based on X-ray tomographic data processed in Drishti v2.4. **A** Dorsal view of the first right trunk appendage. **B** Dorsal view of the third left trunk appendage, red arrows indicate spines on terminal claw of endopods. **C** Dorsal view of the seventh left trunk appendage, red arrows indicate spines on terminal claw of endopods. **D** Ventral view of the eleventh right trunk appendage. **E** Dorsal view of left posterior trunk appendages. Abbreviations: el, exopod lobe; en, endopod; lm, lamella(e); exi, exite(s); hs, head shiled; pd*n*, endopod podomere number; pt, protopod; st, seta(e); tc, terminal claw; T*n*, trunk appendage number; *l*, left; *r*, right. Scalebar: 1 mm in **A**–**E**.

#### Compound eyes

The eyes of *C. eucalla* are compound and of ventral origin. Across all examined specimens of *C. eucalla*, the compound eyes exhibit pronounced positional variability (Figure 2). In specimens preserving both left and right eyes in an essentially symmetrical arrangement, the eyes are situated longitudinally within the anterior one-third to one-half of the cephalic shield (Figures 1 A, B, D; 6 A; Figure 1—figure supplement 1 A, C; Figure 2—figure supplement 2 D, E, F, G). The extent of eye exposure beyond the cephalic margin varies considerably, from nearly the entire eye (Figure 1 B; Figure 1—figure supplement 1 A; Figure 4—figure supplement 4 C) to merely the peripheral rim (Figure 1—figure supplement 1 C; Figure 2—figure supplement 2 E, F, G). Incompletely preserved specimens retaining one eye exhibit comparable positioning within the anterior one-third to one-half of the shield, likewise varying from largely exposed (Figure 7 A; Figure 1—figure supplement 1 B; Figure 2—figure supplement 2 B, C) to only marginally protruding beyond the cephalic margin (Figures 3 A). Because most specimens are dorsoventrally compressed and preserve neither the complete stalk nor its articulation, this positional variation cannot distinguish ocular mobility in life from taphonomic displacement. We therefore treat it as a preservational and morphological observation rather than direct evidence of active protraction, retraction, or antero-posterior movement.

#### Anterior sclerite and hypostome

There is a small sclerite located in front of the hypostome (Figures 3 E, E1, E2; 5 A, B; supplementary video 1, 2, 3). It is narrower and shorter than the hypostome. The hypostome is oval, and covers the anterior gut (Figures 1 B, C, D; 6 A; Figure 1—figure supplement 1 A, C). There is a putative mouth near the posterior margin of the hypostome (Figure 5 A, B).

**Figure 5.**
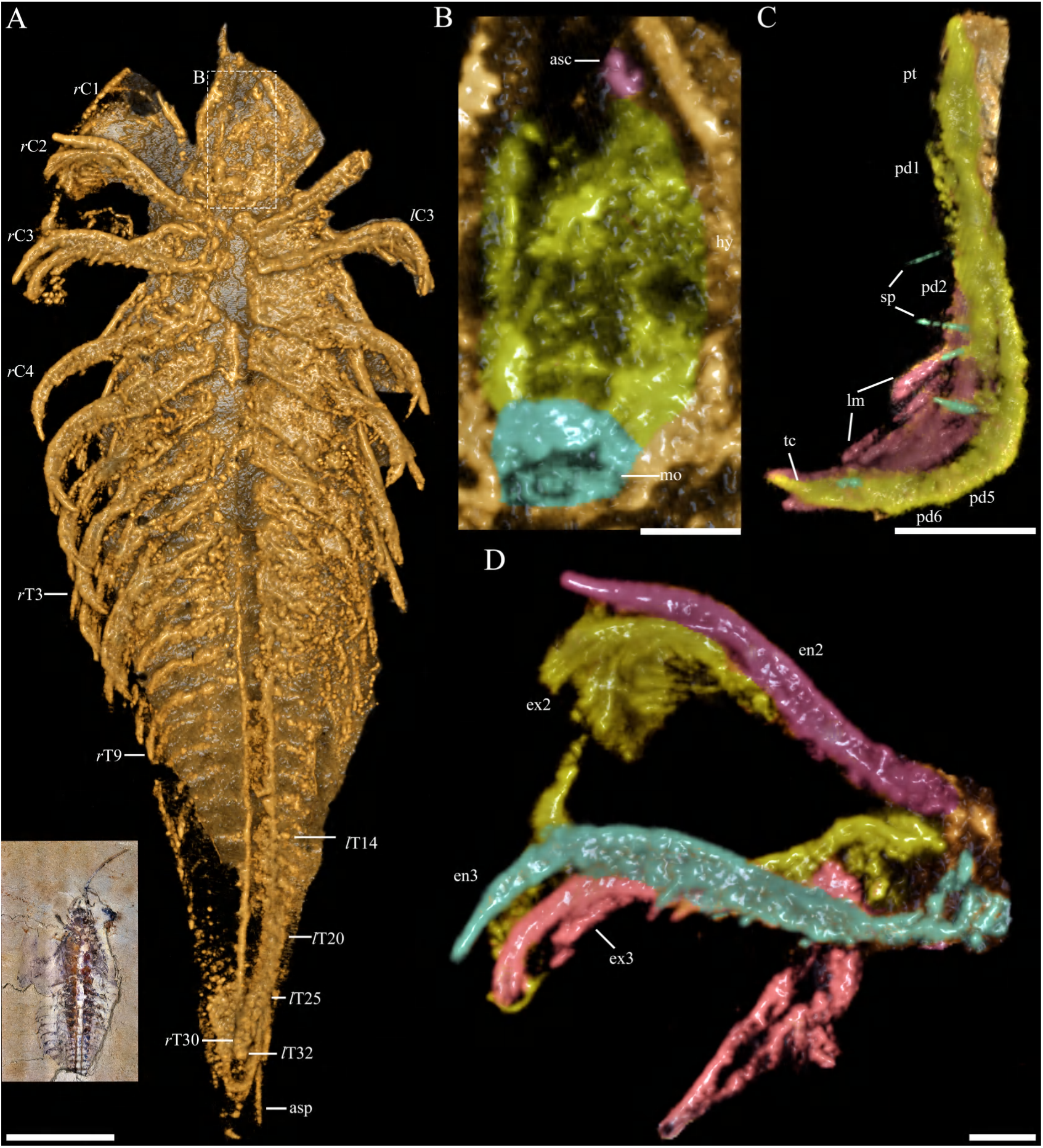
Digital anatomy of *C. eucalla* (YKLP 17370) based on X-ray tomographic data processed in Drishti v2.4. **A** Overview, ventral, with dorsal exoskeleton partially removed. **B** anterior structures beneath the head shield within the dashed box in **A**, with anterior sclerite, hypostome and mouth painted in different colours respectively. **C** Ventral view of the third left cephalic appendage. **D** Ventral view of the second and third right cephalic appendages, with the exopods and endopods painted in different colours. Abbreviations: ant, antenna(e); asc, anterior sclerite; asp, axial spine(s); C*n*, post-antennal cephalic appendage number; en, endopod; ex, exopod; hy, hypostome; lm, lamella(e); mo, mouth opening; pd*n*, endopod podomere number; pt, protopod; sp, spine(s); tc, terminal claw; T*n*, trunk appendage number;*l*, left; *r*, right. Scalebar: 2 mm in **A** and **B**, 1 mm in **C**, 0.5 mm in **D**.

#### Gut and diverticulae

The gut extends from the hypostome to the terminal tergite (Figures 1 B, C, D; 5 A, B; 6 A; Figure 1—figure supplement 1; Figure 2—figure supplement 2 B, D–F), and in YKLP 17363a, about 40% of the hypostome’s length is covered dorsally by the gut (Figure 1 B, C). The diverticulae resembles to those of *Naraoia spinosa* (Zhang et al., 2007; Zhai et al., 2019), but the branch pattern is simpler (Figures 1 B; 6 A; Figure 1—figure supplement 1 A, B).

#### Cephalic appendages

The multi-annulated antenna is thin and long, with over 30 articles (Figures 1 A; 3 A, B, D; 7 A; 8 A, B; Figure 1—figure supplement 1; Figure 2—figure supplement 2 C, F, G; Figure 4—figure supplement 4 C). The articles decrease slowly in length and width from the proximal to distal (Figures 1 A; 3 B; Figure 1—figure supplement 1 B, C; Figure 2—figure supplement 2 C, F, G). An antennal scale attaches at the rear end of the first annula or the base of the second one, extending laterally from probably the inner side (Figure 3 B). Behind the antennae, there are four pairs of appendages that are morphologically similar to the trunk appendages, located adjacent to the anterior margin of the first trunk tergite (Figures 1 A, B, D; 3 A, C). The appendages are biramous, with each consisting of a protopod, endopod, and exopod. They are morphologically similar, and their size progressively increases posteriorly (Figures 3 D; 5 A, D; 6 A, B; 8 B). The morphology of cephalic appendages is similar to these of trunk, especially the anterior trunk appendages (Figures 3 D; 5 A, C, D; 6 B; 8 A, B). Cephalic appendages are small compared to the anterior trunk appendages, tapering anteriorly (Figures 3 D; 5 A, D; 6 B; 8 A, B). The last cephalic appendage pair is smaller than the first trunk appendage pair (Figures 3 D; 5 A; 6 B; 8 B).

#### Trunk appendages

All trunk appendages are biramous and share a similar morphology. Each appendage comprises a protopod with gnathobasic bulges, a six-podomere endopod armed with spines on each podomere, an exopod with a paddle-shaped shaft bearing over 20 lamellae, and a rod-like exite (Figures 3 D; 4; 5 C, D; 6 B, C; 8 C, D; Figure 6—figure supplement 6 B, D). Each of the first six anterior trunk tergites beneath the posterior extension of the head shield covers a pair of biramous appendages (Figures 1 A, B, D; 3 A, C; 6 B). The first or second trunk appendage pair is the largest among all trunk appendage pairs, and the following three or more pairs are comparable in size (Figures 3 D; 4; 5 A; 6 B). All trunk appendages taper posteriorly, and only limb buds could be observed underneath the last several tergites (Figures 3 D; 4 E; 5 A; 6 B; 8 B, F). Behind the sixth trunk tergite, *C. eucalla* shows a decoupling of tergites and appendage pairs (Figures 3 D; 6 A, B; 8 B). Tergites posterior to the sixth trunk segment cover an increasing number of appendage pairs, lacking correspondence between tergite boundaries and limb boundaries (Figures 3 D; 6 A, B; 8 A, B; Figure 1—figure supplement 1 C; Figure 4—figure supplement 4 A, C; Figure 6—figure supplement 6). Variation is observed among different specimens (Figures 3 D; 5 A; 6 A, B; Figure 1—figure supplement 1 C; Figure 4—figure supplement 4 A, C; Figure 6—figure supplement 6; and Figures 5 and 9 in Ramsköld et al., 1997).

**Figure 6.**
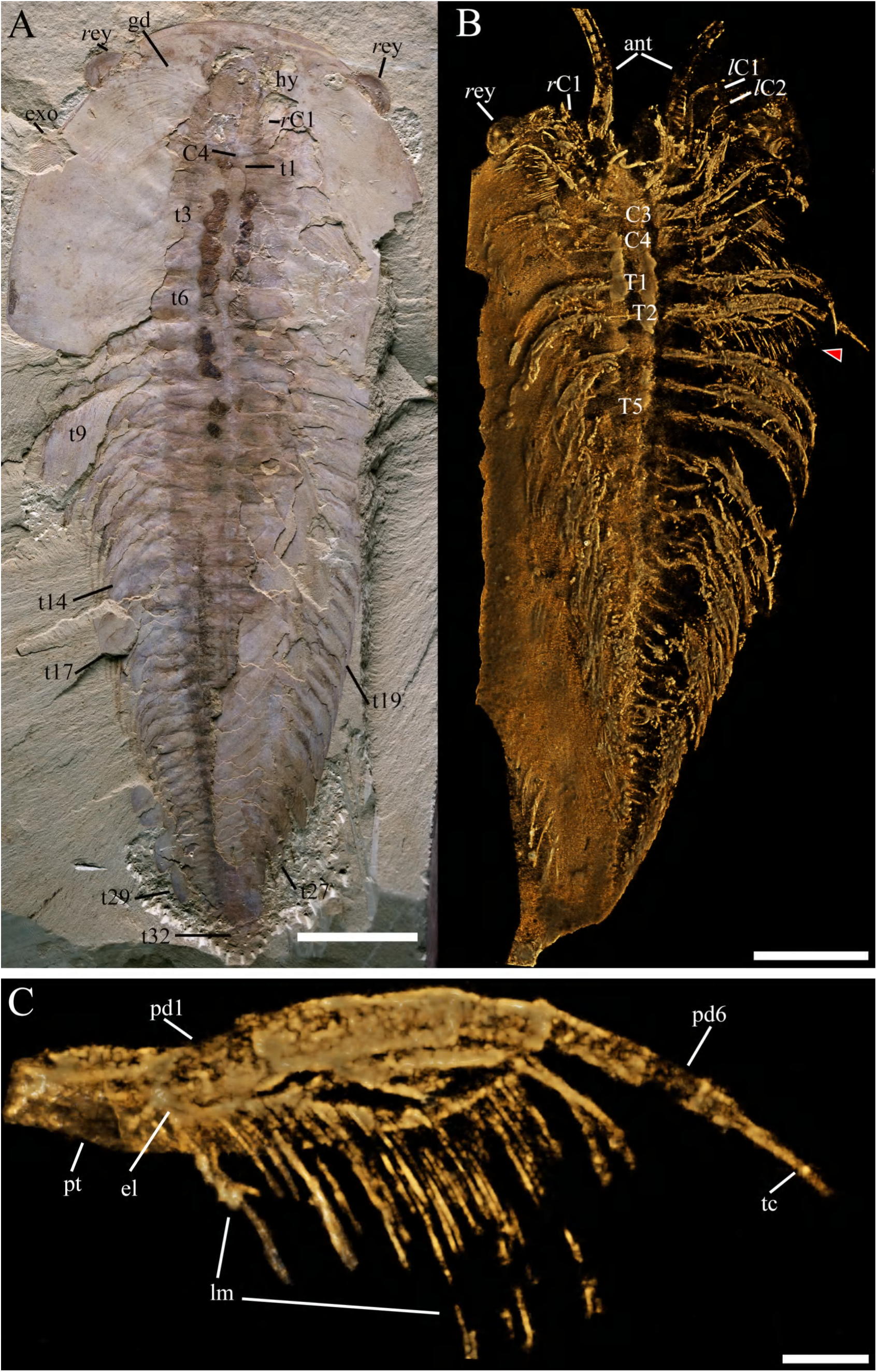
Macro-photographic picture (A) and results from X-ray tomographic data processed in Drishti v2.4 (B, C), YKLP 17361. A Overview, dorsal. **B** Overview, ventral. **C** Ventral view of the second left trunk appendage. Abbreviations: ant, antenna(e); C*n*, post-antennal cephalic appendage number; el, exopod lobe (shaft); ey, eye(s); gd, gut diverticula(e); hy, hypostome; pd*n*, endopod podomere number; pt, protopod; sp, spine(s); tc, terminal claw; T*n*, appendage number; t*n*, tergite number; *l*, left; *r*, right. Scalebar: 1 cm in **A** and **B**, 2 mm in **C**.

### Phylogenetic results

Under the implied-weight parsimony analysis shown in Figure 9 A, all Xandarellida form a robust monophyletic clade anchored by a 100% consensus frequency node, indicating the cohesive morphological synapomorphies uniting this taxon under parsimony criteria. *Zhugeia acuticaudata* is resolved as the earliest-diverging taxon at the base of Xandarellida, succeeded by two derived subclades. One derived subclade unites *L. deletres* and *Luoluilinella rarus*, recovered in 95% of all most-parsimonious trees—this represents the only node lacking full 100% consensus frequency across the tree, and insufficient morphological information is the primary cause. The other subclade unites *X. spectacullum*, *C. eucalla* and *S. lunaris* into a fully congruent monophyletic assemblage with 100% consensus frequency at its basal node. Within this subclade, *X. spectacullum* and *C. eucalla* form a tightly supported sister group anchored by another node bearing a consensus frequency of 100%, whereas *S. lunaris* constitutes the basal lineage diverging prior to the this sister group. Xandarellida and the lineage consisting of the ((*Tremaglaspis vanroyi*, *P. pergamena*), (*Liwia convexa*, *Soomaspis*, *Tariccoia*)) form a polytomous clade (Figure 9 A).

In the Bayesian analysis, xandarellids is recovered with a posterior probability (PP) of 55% (Figure 9 B). The crown node of this clade forms a four-way polytomy comprising four lineages: *X. spectacullum*, *C. eucalla*, (*L. deletres*, *L. rarus*) and (*Z. acuticaudata*, *S. lunaris*). The (*L. deletres*, *L. rarus*) and (*Z. acuticaudata*, *S. lunaris*) each have full internal support (PP = 100%). The node subtending the (*L. deletres*, *L. rarus*) lineage holds PP = 50%, whereas the split leading to (*Z. acuticaudata*, *S. lunaris*) bears PP = 82%.

We therefore regard monophyly as supported by the parsimony analyses but only moderately recovered by the Bayesian analysis, and we do not use either result to make strong claims about deeper artiopodan relationships.

## Discussion

### Dorsal exoskeleton

The exoskeleton of *C. eucallais* divided into a head shield and trunk, with the anterior six trunk tergites being covered by the posterior portion of the head shield (the last tergite is also a small pygidium). The anterior six trunk tergites have small or no pleurae (Figure 1 B; 3 C). Without the application of micro-CT scanning in the present study, these structures would not be clearly discernible (Chen et al., 1996; Ramsköld et al., 1997; Chen, 2004; Hou et al., 2004, 2017). The last tergite is a pygidium in *X. spectaculum*, and the body is divided into a head shield, thorax and pygidium, the head shield may be composed of two fused or articulated tergites (Hou et al., 1991, 1999, 2017). Following the head shield is a small tergite and a series of trunk tergites, mostly preserved in dorsal-ventral compression (Hou et al., 1991, 2004, 2017; Chen et al., 1996; Chen, 2004; Hou and Bergström, 1997). *S. lunaris* shows a similar dorsal exoskeleton to *X. spectaculum* (Chen et al., 2019; Schmidt et al., 2021). At least two thoracic tergites of *Z. acuticaudata* are covered by its head shield shield (Zhu et al., 2023). In both *L. ramus* (Zhang et al., 2012) and *L. deletres* (Hou et al., 2018), the head shield is relatively broad and short. Notably, a small, discrete tergite is intercalated between the head shield and the succeeding pleurae-bearing trunk tergites; the terminal tergite may alternatively be interpreted as a diminutive pygidium in *C. eucalla*. In addition, a (or more) small tergite(s) situated immediately posterior to the head shield has been documented in three artiopodans: *Zhiwenia coronata* (Du et al., 2019), *Tremaglaspis vanroyi* (Lerosey-Aubril et al., 2013), and *P. pergamena* (Ivantsov, 1999).

The small tergite and pygidium exhibit considerable homoplasy, indicating repeated independent evolutionary transitions rather than shared ancestry (Figure 7—figure supplement 7 D, H). The small tergite(s) (reduced anterior tergites; three steps, CI = 0.333, RI = 0.750, score = 0.286) requires three independent state changes, suggesting that the structural integration between the head shield and anterior tergite is developmentally labile and prone to parallel modification or loss. The pygidium (four steps, CI = 0.250, RI = 0.850, score = 0.167) shows even more extensive homoplasy; its low character score and high step number imply multiple independent origins or reversals across the tree. The relatively high RI of both characters indicates localized lineage conservatism once these structures are acquired, but their scattered distributions overall reflect developmental plasticity and/or convergent functional adaptation rather than reliable phylogenetic signal. Consequently, neither character can be treated as a synapomorphy for any major clade.

**Figure 7.**
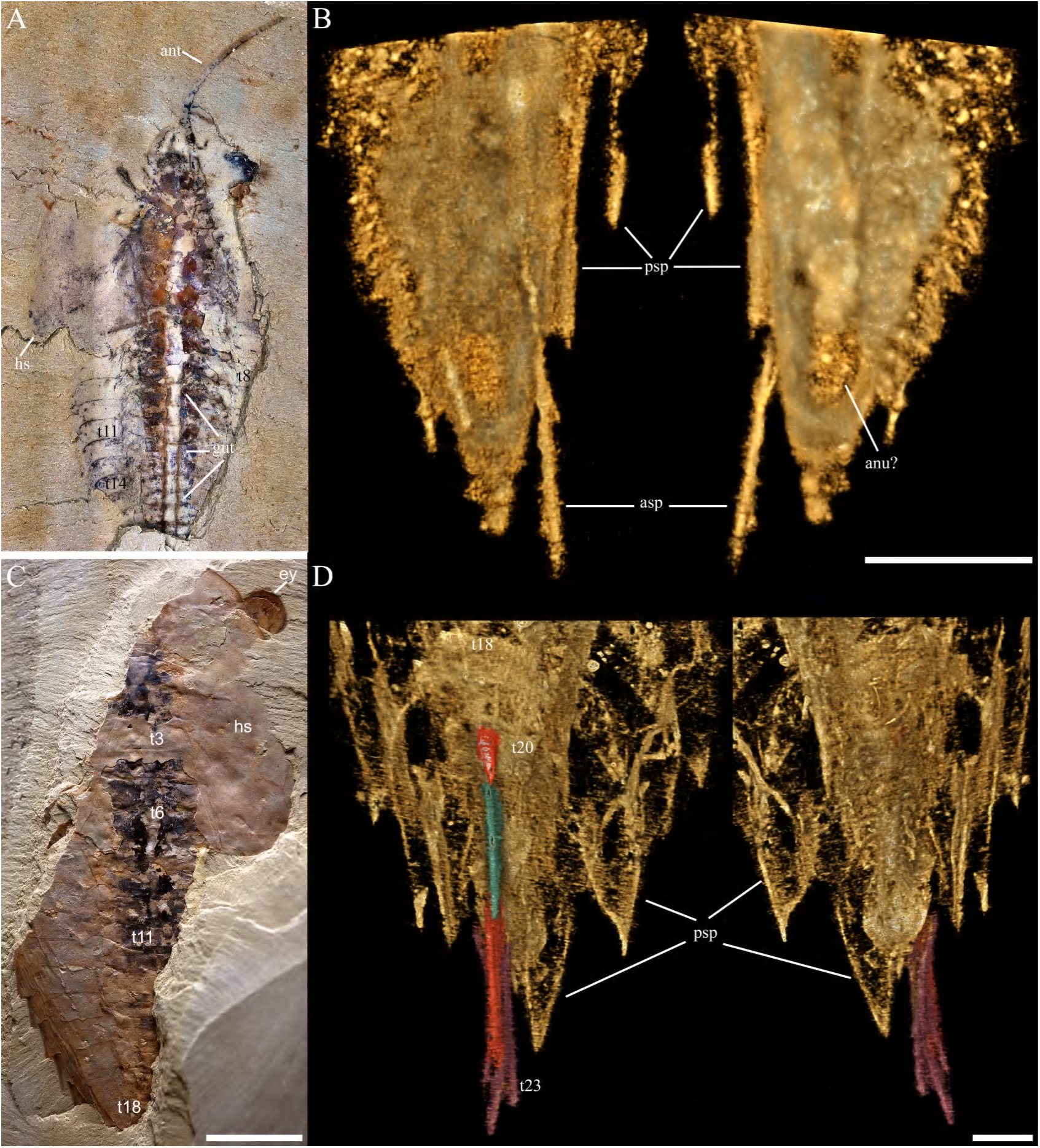
Macro-photographic picture (A, C) and results from X-ray tomographic data processed in Drishti v2.4 (B, D). **A** Overview of YKLP 17370, dorsal view. **B** Dorsal (left) and ventral (right) views of the posterior trunk in **A** hide in the rock. **C** Overview of YKLP 17371, dorsal view. **D** Dorsal (left) and ventral (right) views of posterior trunk in **C** hide in the rock. Abbreviations: ant, antenna(e); anu?, putative anus; asp, axial spine(s); C*n*, post-antennal cephalic appendage number; ey, eye(s); gut, gut content/trace; hs, head shield; psp, pleural spine(s); t*n*, tergite number. Scalebar: 1 cm in **C**, 2 mm in **A**, 1 mm in **B** and **D**.

### Eye slits and exoskeletal bulges

*C. eucalla* lacks the eye slits present in *X. spectaculum* (Hou et al., 1991; Hou and Bergström, 1997; Hou et al., 2017), as well as the lateral notches found in *L. ramus* (Zhang et al., 2012) and *L. deletres* (Hou et al., 2018). *X. spectaculum* has distinct eye slits extending from the eyes to the lateral margin of the head shield, dividing it into two parts (Hou et al., 1991; Hou and Bergström, 1997; Chen, 2004; Hou et al., 1999, 2004, 2017). *P. pergamena* is considered to have eye slits (open sutures in Ivantsov (1999)) that run across the segments transversely. Chen et al. (2019) suggested that the notches found within xandarellid species (*Cindarella*, *Luohuilinella*, *Xandarella* and *Phytophilaspis*, *Sinoburius*) reflect a progressive involution of the perforations accommodating the ventral stalked eyes (Chen et al., 2019). However, this trend is not supported by the morphological comparison and phylogenetic position presented herein.

### Compound eyes

The preserved positions of the eyes cannot distinguish biological mobility from taphonomic displacement because most specimens do not preserve a complete eye stalk or its articulation. We therefore do not infer ocular mobility, protraction–retraction, or an anteroposterior range of movement from positional variability alone.

*C. eucalla*’s eyes are particularly prominent within Artiopoda. The largest compound eyes of *C. eucalla* reach 5.36 mm in an incomplete specimen (YKLP 17367). Stalked compound eyes of *X. spectaculum* (Hou and Bergström, 1997) protrude from the dorsal exoskeletal bulges of the head shield, indicating a limited range of motion within the eye bulges. *X. spectaculum* has smaller eyes than *C. eucalla* when their body length is comparable. Zhao et al. (2013) estimated that *C. eucalla* has more than 2000 ommatidia (the maximum diameter of the measured individual’s eye is about 3.8 mm). *X. spectaculum* has a pair of eyes in middle size compared to *C. eucalla*, *L. deletres* (Hou et al., 2018), *Z. acuticaudata* (Zhu et al., 2023) and *S. lunaris* (Chen et al., 2019; Schmidt et al., 2021). *L. deletres* has a pair of ventral stalked eyes, with a maximum eye diameter of 4.45 mm in a 84.96 mm-long specimen (YKLP 11120) (Hou et al., 2018).

The available material permits a morphometric comparison of relative eye size among xandarellids, but not a direct comparison of visual performance. Stalked compound eyes occur in diverse euarthropods, e.g. *Fuxianhuia proensa* (Ma et al., 2012), *Pygmaclypeatus daziensis* (Schmidt et al., 2022, 2025), *Stanleycaris hirpex* (Moysiuk and Caron, 2022); however, functional inferences in fossils require evidence such as facet number and diameter, interommatidial angles, opticalaxis geometry, or well-preserved stalk articulation (Vannier et al., 2016; Paterson et al., 2020). For example, visual function in the Cambrian crustacean Henningsmoenicaris was inferred from three-dimensionally preserved facets and internal traces, rather than from eye size alone (Schoenemann et al., 2011). Comparable evidence is not available for *C. eucalla*. Accordingly, the ratios in Table 2 are reported as specimen-based morphometric values, not as measures of visual acuity, ecological role, or evolutionary “advancement”.

**Table 1.** Parameters for maximum-parsimony trees and consensus trees built with a 50% frequency cut-off majority-rule criterion, under equal weighting and implied weighting (k = 1, 3, 4, 5, 8, 10, 20).

| Weight | Maximum parsimony trees | consensus tree |
|---|---|---|
| Equal weight | RI=0.707<br>CI=0.387<br>Tree length=279<br>MPT=18<br>Best score=279 | RI=0.700<br>CI=0.382<br>Tree length=283 |
| K=1 | RI=0.670<br>CI=0.359<br>Tree length=301<br>MPT=3<br>Best score=38.695 | RI=0.668<br>CI=0.358<br>Tree length=302 |
| K=3 | RI=0.692<br>CI=0.375<br>Tree length=288<br>MPT=8<br>Best score=25.421 | RI=0.687<br>CI=0.371<br>Tree length=291 |
| K=4 | RI=0.699<br>CI=0.380<br>Tree length=284<br>MPT=19<br>Best score=21.840 | RI=0.685<br>CI=0.370<br>Tree length=292 |
| K=5 | RI=0.705<br>CI=0.386<br>Tree length=280<br>MPT=16<br>Best score=19.166 | RI=0.700<br>CI=0.382<br>Tree length=283 |
| K=8 | RI=0.705<br>CI=0.386<br>Tree length=280<br>MPT=18<br>Best score=14.110 | RI=0.700<br>CI=0.382<br>Tree length=283 |
| K=10 | RI=0.705<br>CI=0.386<br>Tree length=280<br>MPT=18<br>Best score=12.035 | RI=0.700<br>CI=0.382<br>Tree length=283 |
| K=20 | RI=0.700<br>CI=0.386<br>Tree length=283<br>MPT=12<br>Best score=6.997 | RI=0.700<br>CI=0.382<br>Tree length=283 |

Parsimony analyses were run in TNT v1.6 under New Technology Search, combining 50 random addition sequences with ratchet, drift, and tree-fusing heuristics options activated with standard settings, random seed was fixed at 1234, trees up to 10 steps longer than the optimal tree were retained (fit difference = 0.05). Abbreviations: MPT, number of most parsimony trees; RI, retention index; CI, consistency index.

**Table 2.** Maximum measured eye-length/body-length ratios in selected xandarellid specimens.

| Species | Specimen No. | Body<br>Length<br>(mm) | Eye<br>Length<br>(mm) | Eye/Body<br>Ratio<br>(%) | References |
|---|---|---|---|---|---|
| *L. deletres* | YKLP 11120a | 99.07 | 4.73 | 4.77 | Hou et al., 2018, Fig. 5a |
| *Z. acuticaudata* | NIGP 200049 | 46.34 | 2.54 | 5.48 | Zhu et al., 2023, Fig. 1A2 |
| *X. spectaculum* | CN 115285 | 52.81 | 2.43 | 4.60 | Hou et al., 1997, Fig. 71A |
| *S. lunaris* | YRCP 0011 | 7.45 | 0.54 | 7.25 | Chen et al., 2019, Fig. 4b |
| *C. eucalla* | NIGPAS 158641 | 57.00* | 3.80 | 6.67 | Zhao et al., 2013, Fig. 1a |

\* Estimated body length. All eye measurements represent maximum length.

### Anterior sclerite

The Anterior sclerite has been definitively documented only in *C. eucalla* within Xandarellida. In *S. lunaris*, the elongate hypostome appears to comprise an inverted, triangular heart-shaped component and an oval structure, although no detailed evidence currently supports the interpretation that this represents a fused anterior sclerite and labrum (Chen et al., 2019; Schmidt et al., 2021). A discrete sclerite occurs anterior to the hypostome in the interpreted ventral anatomy of *C. eucalla* (Figures 3 E, 5 B; Supplementary Video 1, 2, 3). We refer to it descriptively as an anterior sclerite. Anterior or ocular sclerites reported in other euarthropods differ in position, morphology and preservation, and their homology has not been tested here (Ortega-Hernández, 2015; Zeng et al., 2018, 2020; Park, 2023). The observed association in *C. eucalla* should therefore not be used to infer homology with anterior sclerites of radiodonts (*Lyrarapax unguispinus* (Ortega-Hernández, 2015), *Anomalocaris saron* (Ortega-Hernández, 2015), *Anomalocaris canadensis* (Daley and Edge-combe, 2014), *Hurdia victoria* and *Amplectobelua symbrachiata* (Zeng et al., 2018)), megacheirans Megacheira (*Kootenichela deppi*, *Worthenella cambria*, *Jianfengia multisegmentalis*) (Legg et al., 2013; Aria et al., 2021; Zhang et al., 2022b), fuxianhuiids (*Fuxianhuia protensa*, *Fuxianhuia xiaoshibaensis*, *Apankura machu*, *Alacaris mirabilis*, *Chengjiangocaris longiformis*, *Xiaocaris luoi*, *Guangweicaris spinatus*) or other artiopodans (*Misszhouia longicaudata*, *Helmetia expanse* (Ortega-Hernández et al., 2013; Zeng et al., 2020), *Kuamaia lata*) (Budd, 2008; Ortega-Hernández, 2015). The present result documents a candidate anterior sclerite in *C. eucalla* and identifies a character requiring broader comparative study.

### Natant hypostome and gut

The hypostome plays a pivotal role in the evolution, ecology, and classification of early arthropods (Scholtz and Edgecombe, 2005; Liu et al., 2016; Ortega-Hernández et al., 2017b, 2019; Liu et al., 2020; Budd, 2021; El Albani et al., 2024). Three types of hypostome were identified in artiopods: natant, conterminant, and impendent (Fortey, 1990). Among the xandarellids, *C. eucalla*, *X. spectaculum* (Hou et al., 1991; Hou and Bergström, 1997; Hou et al., 2017) and *S. lunaris* (Hou et al., 1991; Chen et al., 2019; Schmidt et al., 2021) have typical natant hypostomes (Figure 1 C, D). *C. eucalla*’s hypostome is oval in shape, and its midgut extends from the posterior third of the hypostome to the rear of the body (Figure 1 C, D), indicating that the midgut opening is located under the hypostome, while the mouth opening is likely at the end of the gut margin covered by the hypostome. *X. spectaculum* has a subelliptic hypostome (Hou and Bergström, 1997). while *S. lunaris* has a natant hypostome with a sharp anterior end and a long, oval-shaped posterior end (Chen et al., 2019; Schmidt et al., 2021). The hypostome of *C. eucalla* is oval in the available material. A gut trace extends posteriorly from the cephalic region to the posterior trunk (Figures 1 C,D). Dorsoventral compression and the absence of a directly resolved mouth–labrum complex prevent a confident reconstruction of the exact mouth position or hypostome attachment. The natant configuration is therefore recorded as morphology only. It does not, by itself, test feeding mode or ecological niche; no niche-partitioning inference is made here.

### Antennae and antennal scale

The antennae of Artiopodans are typically uniramous, long, and multi-segmented, with some species possessing setae on the segments (Hou et al., 2017; Zeng et al., 2017; Lerosey-Aubril and Ortega-Hernández, 2019; Holmes et al., 2020; Zhang et al., 2021; Jiao et al., 2022). The antennae of *C. eucalla* and *S. lunaris* (Chen et al., 2019; Schmidt et al., 2021) differ from other Artiopodans, such as *Eoredlichia intermedia* (Hou et al., 2017) and *Retifacies abnormalis* (Zhang et al., 2022a), in terms of length and antennal scales at the base. Antennal scales in Xandarellida vary interspecifically in segmental position and number. *C. eucalla* has a single scale on the second antennal segment, whereas *S. lunaris* (Chen et al., 2019; Schmidt et al., 2021) bears one scale on either the second (YKLP 11407) or sixth segment (YRCP 0011, Hz-f-10-45) (Chen et al., 2019; Schmidt et al., 2021). Although sensory antennal structures are widespread among extant arthropods (Actias luna, Antheraea polyphemus and Bombyx mori in Vogel (1983); Bombus terrestris in Spaethe et al. (2007); Hemileuca eglanterina and Cydia pomonella in Symonds et al. (2012); Uraba lugens in Johnson et al. (2017)), any mechanoreceptive or chemoreceptive function for these scales remains speculative in the absence of preserved neuroanatomy. Prominent antennal bristles are absent in *C. eucalla*, *X. spectaculum* (Hou et al., 2017), *S. lunaris* and *L. deletres* (Hou et al., 2018); however, the hypothesis that scales functionally compensate for this absence is unsupported by the available morphological data.

### Post-antennal cephalic appendages

The morphology of post-antennal cephalic appendages is crucial for studies of arthropod evolution, classification, and ecology (Whittington, 1980; Briggs and Whittington, 1997; Wills, 1998; Waloszek et al., 2005; Lamsdell et al., 2013; Yang et al., 2013; Du et al., 2023). Cambrian euarthropods’ post-antennal cephalic appendages suggest a close connection to feeding (Hou and Bergström, 1997; Waloszek et al., 2007; Hou et al., 2017). *L. deletres*, *C. eucalla*, *S. lunaris* and *X. spectaculum* have 3, 4, 4, and 6 post-antennal cephalic appendage pairs, respectively (Hou et al., 2017, 2018; Schmidt et al., 2021). The post-antennal cephalic appendages of these four species show a trend of increasing size from anterior to posterior, and show morphological differentiation in *S. lunaris*(exopod and endopod podomere number varied towards the posterior of the body). The cephalic and trunk appendages of *X. spectaculum* (Hou and Bergström, 1997; Hou et al., 2017) and *L. deletres* (Hou et al., 2018) remain poorly resolved. The cephalic appendages of *S. lunaris* possess highly segmented endopods and exopods but lack endites, whereas the corresponding appendages of *C. eucalla* closely resemble those of *Triarthrus eatoni* (Hou et al., 2023): each endopodal podomere of the post-antennal appendages carries clustered spines, with terminally divided into multiple terminal branches.

The post-antennal cephalic appendages of *C. eucalla* and *T. eatoni* (Hou et al., 2023) share several gross features, including spiny endopodal podomeres. This resemblance is a descriptive comparison and does not independently establish feeding mode. A functional inference would require a defined mechanical hypothesis and evidence for relevant joint mobility, cuticular strength, muscle arrangement, wear, or biomechanical performance, none of which is available here.

### Trunk appendages

#### Trunk limbs

*C. eucalla* has biramous appendages consisting of a protopod, an endopod with six podomeres and clusters of spines, an exopod with a shaft fringed by lamellae and a distal large lamella with marginal setae (Figures 3 D; 4; 5 A, C, D; 6 C; 8 B, C, D; Figure 4—figure supplement 4 5 B, D in this study; Figures 4, 5, 8, 9–14 in Ramsköld et al. (1997)). Trunk appendages in some trilobites possess gnathobase at their base, an endopod with clusters of spines on its podomeres, and a forked terminal claw (Whittington, 1980; Fortey and Owens, 1999; Fortey, 2004, 2014). The gnathobase of *C. eucalla* appears to be less robust than that of *Limulus polyphemus* or *Sidneyia inexpectans* (Bicknell et al., 2018, 2021). Differences in the trunk endopod of *X. spectaculum* (Hou et al., 1991; Hou and Bergström, 1997; Hou et al., 2017), *C. eucalla*, and *L. deletres* (Hou et al., 2018) mainly involve the number of podomeres, the presence of endites and gnathobase. Several studies suggest that these flap-like structures on exopod of horseshoe crabs can be used for respiration (Gray, 1957; Henry et al., 1996; Suzuki et al., 2008). The trunk endopods of *C. eucalla* and *L. deletres* (Hou et al., 2018) are relatively robust, and no distinct gnathobase has been observed in available material, but there are clusters or pairs of spines on the endopod segments (Figure 3 D; 4 D; 5 B, C; 6 B). Each endopod podomere of *C. eucalla* possesses a cluster of spines, while each endopod podomere of *L. deletres* may bear one or a pair of sharp spines, and no gnathobase or spines on endopod podomeres have been observed in *S. lunaris* (Hou et al., 2018; Chen et al., 2019). In terms of the composition and morphology of the trunk appendages, taxa *C. eucalla*, *X. spectaculum* (Hou and Bergström, 1997; Hou et al., 2017), *S. lunaris* (Chen et al., 2019; Schmidt et al., 2021) and *L. deletres* (Hou et al., 2018) exhibit comparable yet non-identical appendage configurations. The morphological organization of the trunk appendages in *C. eucalla* closely resembles that of *T. eatoni* (Hou et al., 2023), implying a broadly comparable habitat. From a functional morphological perspective, the trunk appendages of *C. eucall* a further show marked similarity to those of *Limulus polyphemus* and *Sidneyia inexpectans* (Bicknell et al., 2018, 2021), indicating possible convergent capabilities.

#### Exite

An exite is a lateral outgrowth associated with the limb base (Liu et al., 2021). In *C. eucalla*, a rod-like lateral structure is interpreted as an exite on positional and morphological grounds (Figures 3 D; 4 E). Previous microtomographic work inferred homology between lamellar exites in some megacheirans (e.g. *Leanchoilia illecebrosa*, *Leanchoilia obesa*) and artiopodans (e.g. *Naraoia spinosa*, and *Retifacies abnormalis*) (Liu et al., 2021). That comparative hypothesis does not by itself establish homology for the rod-like structure in *C. eucalla* or an ancestral state for all major arthropod lineages. We therefore code and discuss the structure as a candidate exite, with its homology left open.

### Segmental mismatch

In early arthropods with multi-segmented thorax composed of both dorsal and ventral elements (generally for limb pairs)—such as leachioliids (Liu et al., 2016), numerous artiopodans excluding xandarellids (Hou et al., 2017) and some trilobites (Hughes, 2003); and in extant arthropods including Chilopoda, chelicerates and hexapods, the condition is a one-to-one correspondence between ventral and dorsal segments (Minelli et al., 2013). Nevertheless, in some arthropod taxa, dorsal and ventral segment counts do not match; this condition is termed segmental mismatch (Minelli et al., 2013). The segmental mismatch of *C. eucalla* is remarkably complex. Intraspecifically, the number of trunk tergites varies extensively, with five distinct states observed: 22, 23, 24, 28, and 32 (see dorsal exoskeleton in the Description). In *C. eucalla*, the correspondence between posterior tergites and limb pairs can be assessed only where both dorsal and ventral structures are sufficiently preserved (Figure 6—figure supplement 6). The available specimens show that posterior tergite boundaries need not coincide with limb-pair boundaries, but incomplete preservation prevents a complete, specimen-by-specimen count for every tergite. We provide all observable correspondences, specimen numbers and confidence assessments in Figure 6—figure supplement 6. Under the present character coding, the mapped distribution of the relevant state is concentrated in xandarellids; this result is not evidence that dorsoventral decoupling is unique to that clade or that it had a particular developmental cause.

### Ecological implications

The new material provides a substantially improved anatomical basis for future ecological tests, but it does not directly establish diet, locomotory performance, speed or habitat. The natant hypostome, compound eyes and biramous appendages of *C. eucalla* (Hou et al., 2023) are compatible with more than one functional interpretation. Although some appendicular features resemble those of *T. eatoni*, *Limulus polyphemus* and *Sidneyia inexpectans* (Bicknell et al., 2018, 2021), these taxa differ in phylogenetic position, body plan and preservational context. We therefore limit the present contribution to morphological comparison and regard feeding, locomotion and habitat as unresolved hypotheses.

### Conclusions

This study elucidated the ventral organization of *C. eucalla*, including its anterior sclerite, appendage morphology, axial spines and segmental mismatch. All xandarellids are robustly recovered as a monophyletic clade under maximum parsimony; recovered, but moderately supported in Bayesian analysis. However, their interrelationships with other artiopodan taxa remain equivocal.

## Methods

### Materials

Seventeen specimens (YKLP 17360–17376) of *C. eucalla* deposited at the Yunnan Key Laboratory for Palaeobiology, Yunnan University were collected in this study. All specimens were collected from the following places: Haikou, Jinning and Anning in Kunming. They stratigraphically belong to the *Eoredlichia-Wutingaspis* biozone of Yu’anshan Member, Chiungchussu Formation (Cambrian Series 2, Stage 3). The remarks on synonymy is based on Hou and Bergström (1997); Ramsköld et al. (1997); Edgecombe and Ramsköld (1999); Hou et al. (2004, 2017).

Seventeen specimens of *C. eucalla* (YKLP 17360–17376), housed in the Yunnan Key Laboratory for Palaeobiology, Yunnan University, were examined in this study. The synonymy assessment is based on direct re-examination of *C. eucalla* and on published descriptions Hou and Bergström (1997); Ramsköld et al. (1997); Edgecombe and Ramsköld (1999); Hou et al. (2004, 2017) and figures of the comparative xandarellid taxa; no new observations of comparative type material are claimed unless stated explicitly.

### Observation and documentation

Macrophotography was captured in Canon EOS 5DS R camera (DS126611, Canon Inc., Tokyo, Japan) with MACRO PHOTO LENS MP-E 65 mm, with directional illumination provided by a LEICA LED5000 MCITM (Leica Microsystems, Wetzlar, Germany) (Figures 1 A, B, D; 6 A; 7 A, C; Figure 1—figure supplement 1; Figure 2—figure supplement 2; Figure 3—figure supplement 3 A, C; Figure 4—figure supplement 4 A, C). Fluorescence microscopic images were photographed with a Leica DFC7000T CCD linked to a Leica M205 FA fluorescent microscope (Leica Microsystems, Wetzlar, Germany) (Figures 1 C; Figure 3—figure supplement 3 B, D; Figure 4—figure supplement 4 B, D). All Images were processed and arranged into figures with Adobe Photoshop CC 2018.

### X-ray computed tomography and 3D rendering

X-ray computed tomography were performed on a Zeiss Xradia 520 Versa (Carl Zeiss X-ray Microscopy, Inc., Pleasanton, CA, USA) at the Yunnan Key Laboratory for Palaeobiology, Institute of Palaeontology, Yunnan University, Kunming, China. Scanning parameters of the four (YKLP 17369, YKLP 17370, YKLP 17361 and YKLP 17371) specimens are given in Table 3. All TIFF stacks were processed with software Drishti (Version 2.4) to enable digital dissections of various structures (Figures 3–5; 6 B, C; 7 B, D).

**Table 3.** X-ray microtomography acquisition parameters for six scans of four individuals of *C. eucalla*. Each row represents one acquisition.

| Collection number | Figures | Pixel size(um) | Detector | Voltage (kV) | Power (W) | Filter | Rotation | Projection | exposure time (s) |
|---|---|---|---|---|---|---|---|---|---|
| YKLP 17369a | Fig.3A-C | 11.35 | FP | 60 | 5 | no | -120°–+120° | 1201X6 | 0.4 |
| YKLP 17369b | Figs. 3D-E2, 4 | 11.61 | FP | 60 | 5 | no | -120°–+120° | 1201X5 | 0.4 |
| YKLP 17370 | Figure 5A | 11.51 | 0.4X | 60 | 5 | no | -103°–+103° | 801x3 | 4.5 |
|  | Figures 5B-D, 7B | 9.48 | 0.4X | 60 | 5 | no | -103°–+103° | 1001x4 | 4 |
| YKLP 17361 | Figure 6B, C | 28.96 | FP | 80 | 7 | no | -117°–+117° | 1201X6 | 0.3 |
| YKLP 17371 | Figure 7D | 33.10 | FP | 80 | 7 | no | -116°–+116° | 1201X5 | 0.4 |

YKLP, abbreviation for Yunnan Key Laboratory for Palaeontology.

### 3D modelling

Complete 3D models of *C. eucalla*(Figure 8) were reconstructed in Blender 2.9 by starting with basic meshes such as spheres and cubes, then refining their shapes using the sculpting tool.

**Figure 8.**
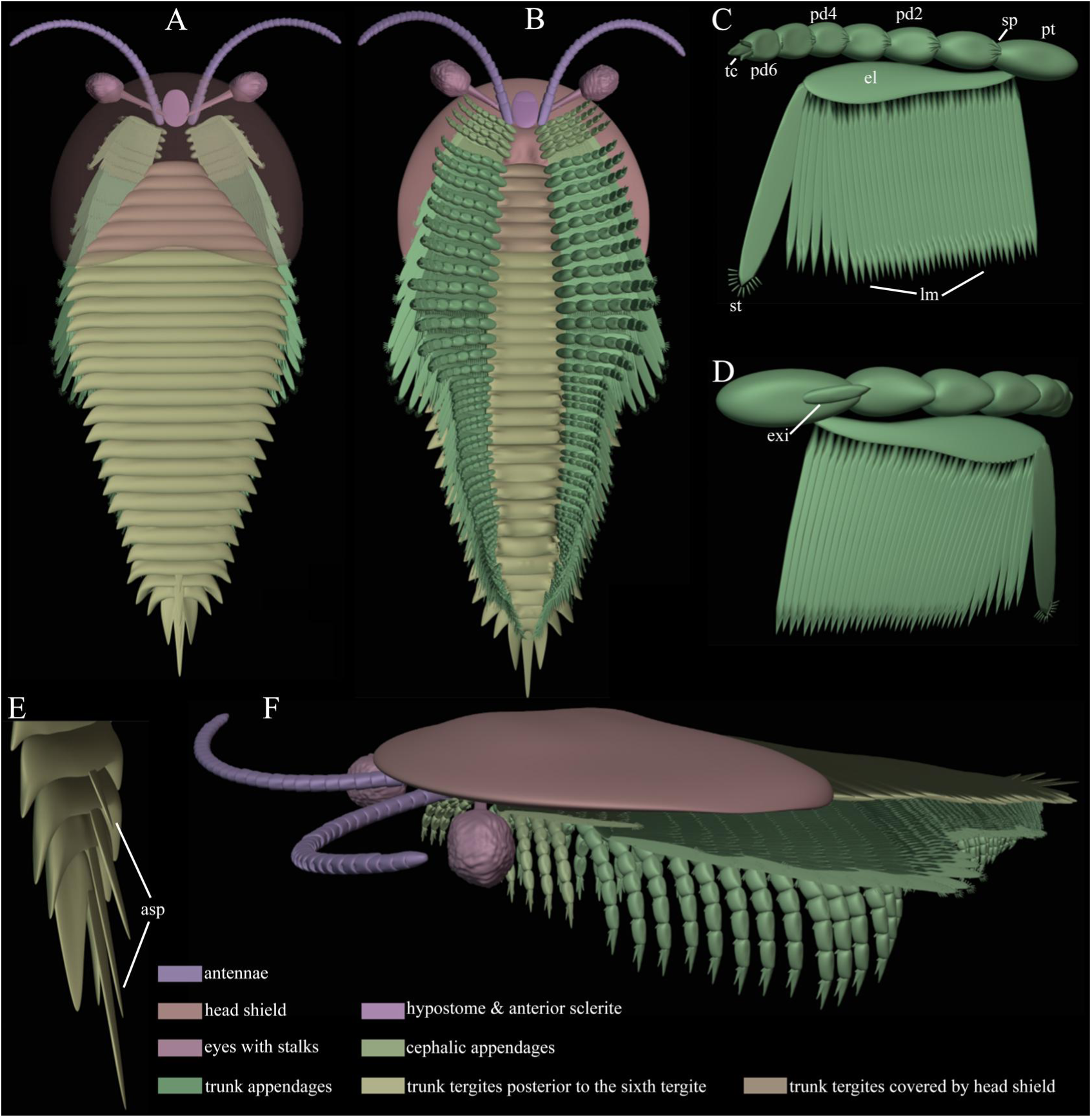
Three-dimensional Blender model of *C. eucalla*. The model is a schematic reconstruction. Eye/stalk pose, dorsoventral thickness, limb cross-section and the volume of the protopod are interpretive where they are not directly resolved by the compressed specimens; the reconstruction should not be read as independent evidence for those features. **A** Overview in dorsal view with the head shield transparent. **B** Overview in ventral view. **C** Biramous appendage in ventral view. **D** Biramous appendage in ventral view. **E** Last four trunk tergites with axial spines in antero-lateral oblique view. **F** Overview in lateral left view. Abbreviations: asp, axial spine(s); el, exopod lobe/shaft; ey, eye(s); exi, exite; lm, lamella(e); pd*n*, podomere number; pt, protopod; sp, spines; st, seta(e); tc, terminal claw. Please note that the three-dimensional reconstruction of the dorsal and ventral aspects incorporates artistic interpretation, as dorsoventral compression of the fossil precludes accurate determination of dorsal thickness and the dorsoventral depth of appendages. Not to scale.

### Phylogenetic analysis

The character matrix employed in the phylogenetic analyses encompasses 70 taxa and 93 characters, as detailed in the character matrix (Morphobank project 6379). This study utilizes an updated iteration of the dataset previously employed by Zhang et al. (2022a), incorporating additional information in this study and five species that have been recently documented by Berks et al. (2023) and Zhu et al. (2023). The morphological characters of species *Acanthomeridion serratum* (Du et al., 2024; Wu et al., 2025; Yang et al., 2026), *Kuamaia lata* (O’Flynn et al., 2025), and *Pygmaclypeatus daziensis* (Schmidt et al., 2025) were updated based on recent studies focusing on these taxa. Parsimony analyses were run in TNT v1.6 under New Technology Search, combining 50 random addition sequences with ratchet, drift, and tree-fusing heuristics options activated with standard settings, random seed was fixed at 1234, trees up to 10 steps longer than the optimal tree were retained (fit difference = 0.05), and duplicate suboptimal trees were discarded (Goloboff, 1999; Goloboff et al., 2008; Goloboff and Catalano, 2016; Goloboff and Morales, 2023).

Equal-weight parsimony was used as a baseline in which all characters contribute equally. Implied weighting was additionally used because it downweights characters that display greater homoplasy during tree search. We explored k = 1, 3, 4, 5, 8, 10 and 20 as a sensitivity analysis, rather than treating any single k value as intrinsically optimal. Figure 9 presents the k = 5 result only as a representative implied-weight analysis; Table 1 reports the placement and support of Xandarellida, *C. eucalla* and *Sidneyia* across all k values and under equal weights. If a focal relationship changes among analyses, we report it as unstable. For Bayesian inference, we use the Mk model, nruns=3, nchains=4, temp=0.1, swapfreq=30, nswaps=2, diagnfreq=5000, printfreq=10000, samplefreq=1000, ngen=50000000, checkpoint=yes, checkfreq=100000, contype=halfcompat, relburnin=yes, burninfrac=0.25), convergence diagnostics ASDSF<0.01. The branch labels show posterior probabilities.

**Figure 9.**
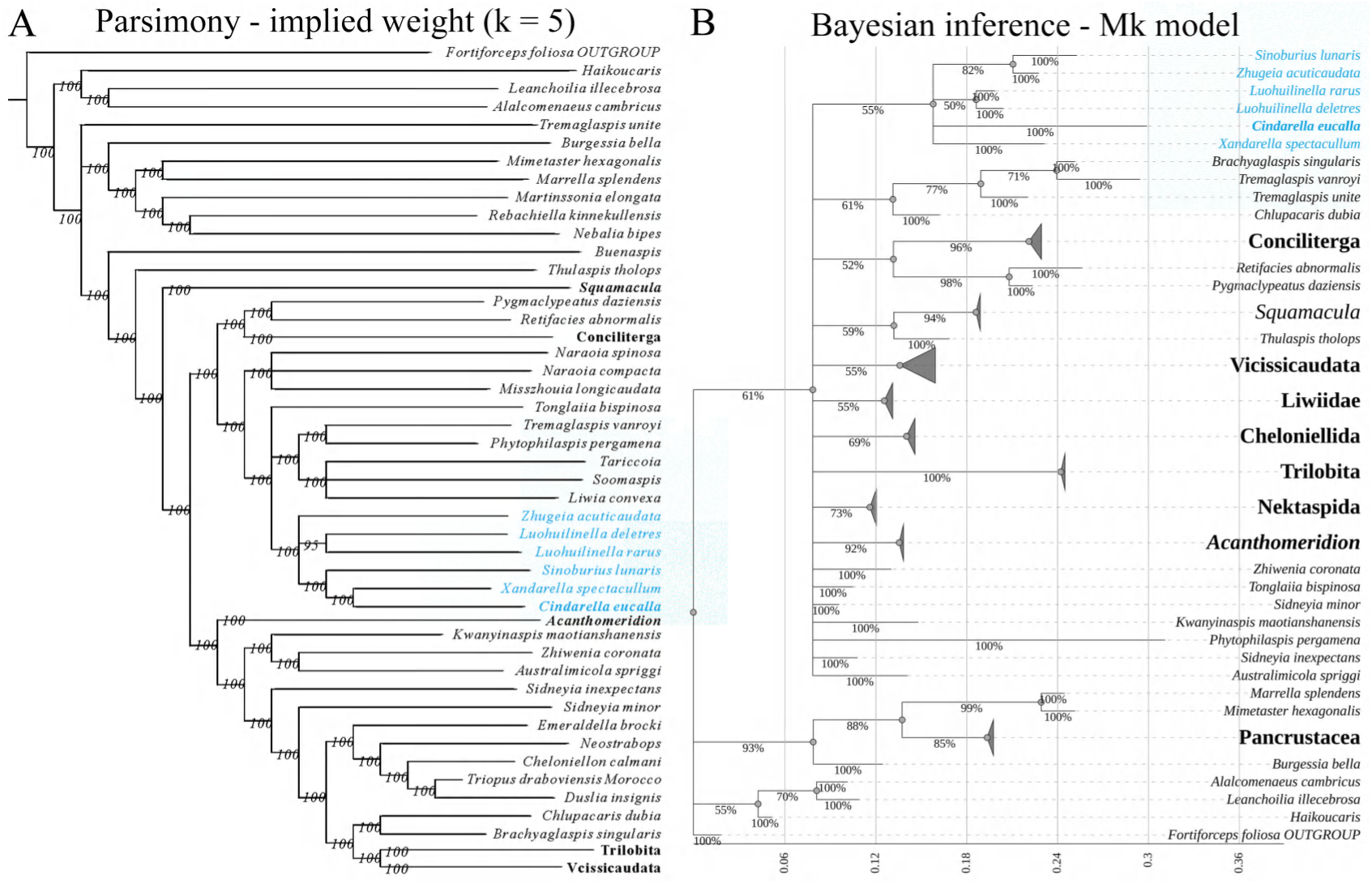
Results of parsimony and Bayesian analyses. Multiple species within the same genus are indicated as a single genus and marked in bold, Xandarellida is highlighted in blue, with *C. eucalla* presented in bold font. **A** Implied weight parsimony, new technology search, 50% cut-off majority-rule consensus tree (k = 5, consistency index (CI) = 0.382, retention index (RI) = 0.700, tree length = 283) computed from the 16 most parsimonious trees, branch labels indicate clade frequencies (>50%) across the set of most parsimonious trees. **B** Bayesian inference under Mk model, branch labels show posterior probabilities (nruns=3, nchains=4, temp=0.1, swapfreq=30, nswaps=2, diagnfreq=5000, printfreq=10000, samplefreq=1000, ngen=50000000, checkpoint=yes, checkfreq=100000, contype=halfcompat, relburnin=yes, burninfrac=0.25).

## Acknowledgments

The comments and suggestions of the editor and three anonymous reviewers greatly improved the manuscript. We extend our heartfelt gratitude to Dayou Zhai (Yunnan University) for his invaluable assistance during the fieldwork phase of our study. We are also grateful to Dr. Javier Ortega-Hernández (Harvard University) for his constructive advice during the preparation of this work.

This study is supported by grants from the Natural Science Foundation of Yunnan Province (202301AS070049; 202401BC070012), National Key R & D Program of China (2025YFF0811700) to Y.L., who is further supported by the Yunnan Revitalization Talent Support Program. M.Z. is supported by a Yunnan University Post-Doctoral Research Fund (W8163003). The funders had no role in study design, data collection and interpretation, or the decision to submit the work for publication.

## Additional information

### Author contributions

Maoyin Zhang, investigation, methodology, software, visualization, formal analysis, writing – original draft, review and editing; Yu Liu, supervision, conceptualization, data curation, formal analysis, funding acquisition, methodology, project administration, resources, validation, visualization, writing, review and editing; Huijuan Mai, software, Visualization; Michel Schmidt, software, visualization; Xianguang Hou, investigation, writing – review and editing.

## Data availability

All figured *C. eucalla* specimens examined and full raw scanning data in this study are permanently deposited in the Yunnan Key Laboratory for Palaeobiology, Institute of Palaeontology, Yunnan University, Kunming, China, with catalogue numbers YKLP 17360–17376. Reconstructed 3D models are available from Michel Schmidt. Supplementary Videos 1–3 are uploaded on elife. Datasets for the phylogenetic analyses are uploaded to Morphobank, Project 6379.

**Video 1. Rotation of the 3D model of YKLP 17369b along the longitudinal body axis.** The 3D reconstruction of YKLP 17369b presented in Figure 3 D is rotated approximately 360° along the anteroposterior body axis, providing a complete view of the overall morphology.

**Video 2. Rotation of the 3D model of YKLP 17369b along the transverse body axis.** The 3D reconstruction of YKLP 17369b presented in Figure 3 D is rotated approximately 360° along the dorsoventral body axis, highlighting lateral and dorsal-ventral morphological features.

**Video 3. Rotation of the ventral hypostome of YKLP 17369b.** The 3D model of YKLP 17369b shown in Figure 3 E is rotated 180° along the anteroposterior axis and 180° along the dorsoventral axis successively, illustrating the relative position of the ventral hypostome from multiple perspectives.

---

**Figure 1—figure supplement 1.**
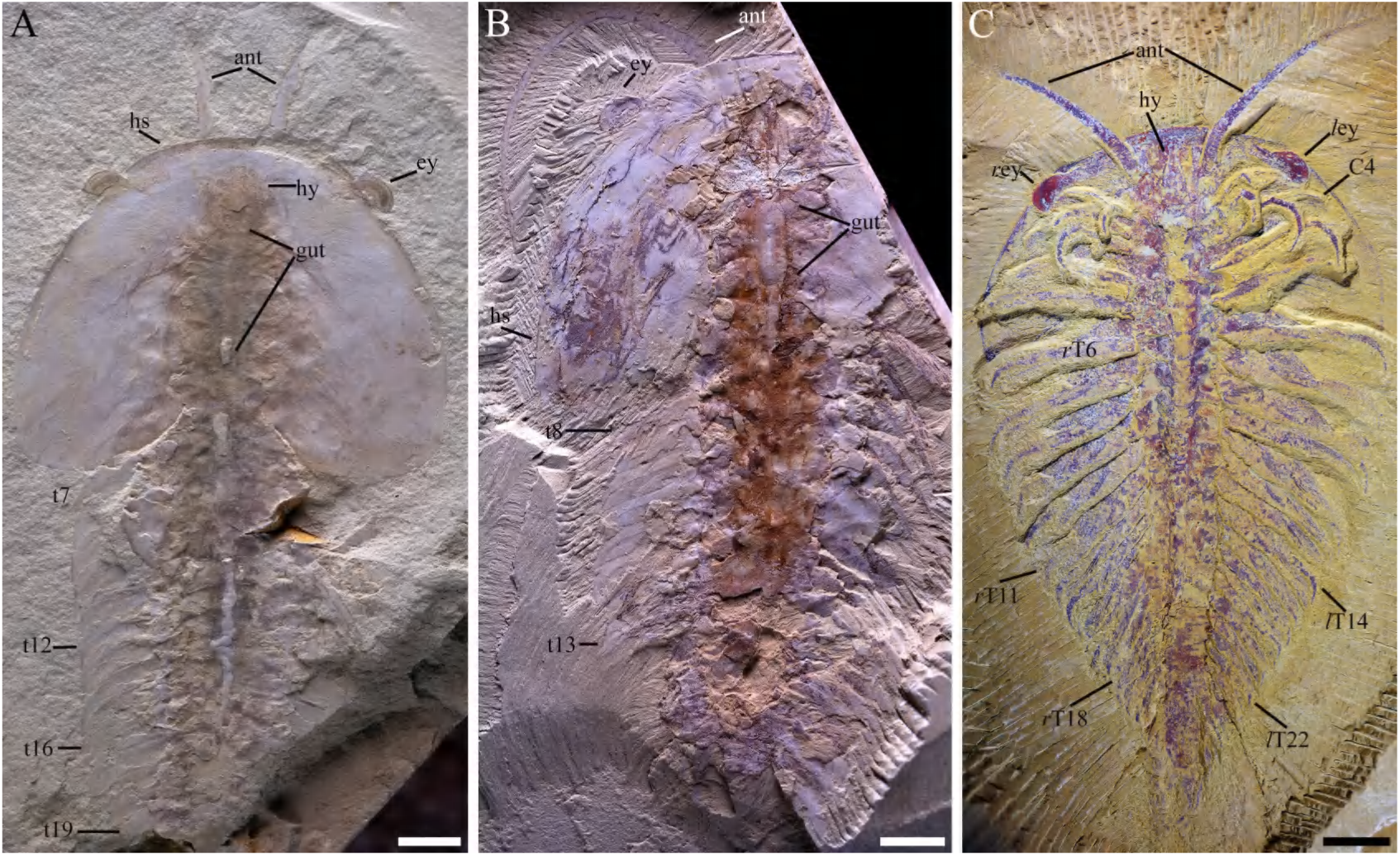
Macro-photographic images of C. eucalla. A Dorsal view of YKLP 17363b. B Dorsal view of YKLP 17364. C Dorsal view of YKLP 17375. Abbreviations: ant, antenna(e); C*n*, post-antenna cephalic appendage number; ey, eye(s); gut, gut content/trace; hs, head shiled; t*n*, tergite number; *r*, right; *l*, left. Scalebar: 5 mm.

**Figure 2—figure supplement 1.**
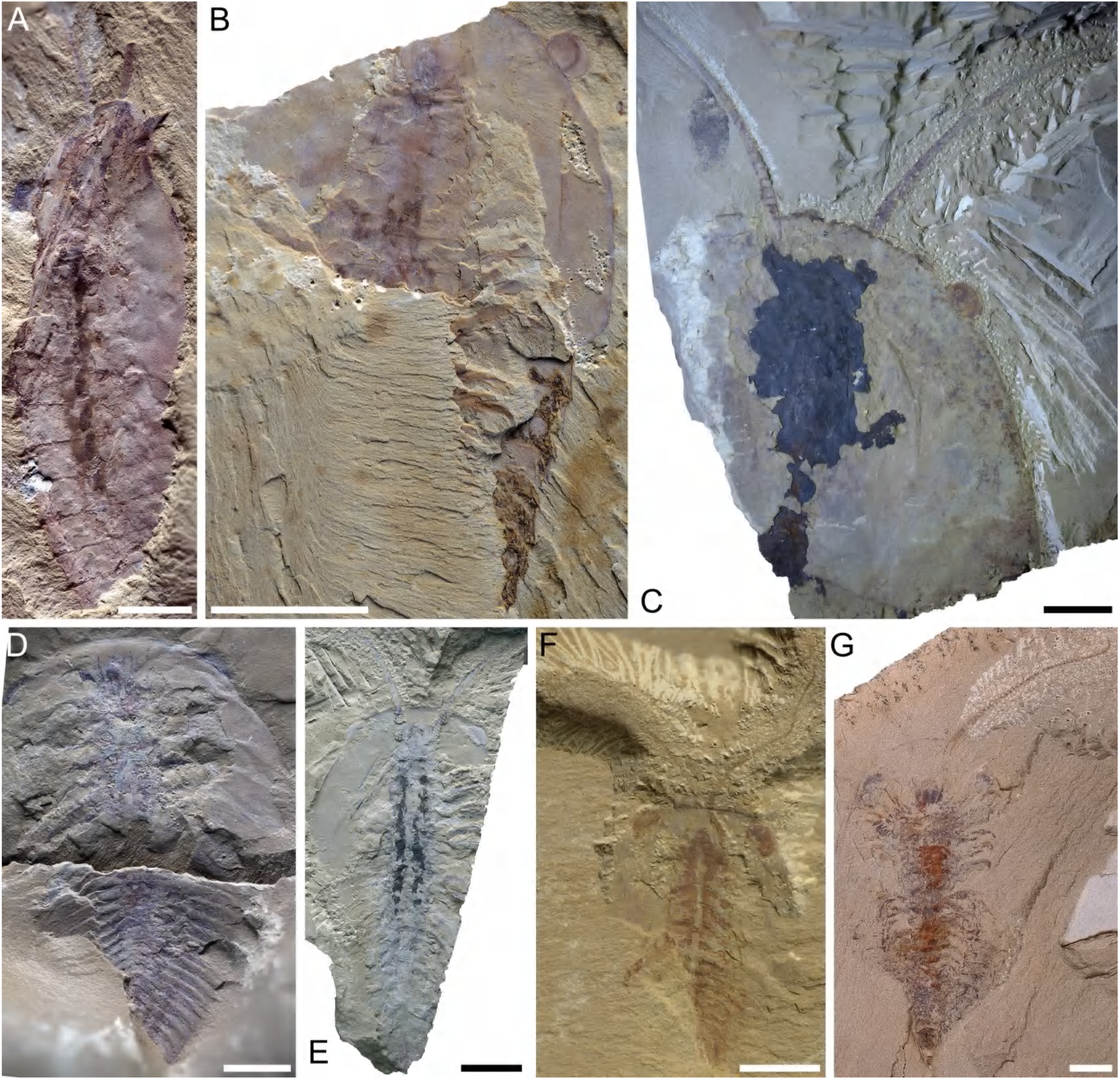
Head shield and eyes of C. eucalla. A YKLP 17366, Dorsal view. B YKLP 17368, dorsal view. C YKLP 17367, dorsal view. D YKLP 17365, dorsal view. E YKLP 17376, dorsal view. F YKLP 17373, dorsal view. G YKLP 17372, dorsal view. Scalebar: 2 mm in G, 5 mm in A, D and F, 1 cm in B, C and E.

**Figure 2—figure supplement 2.**
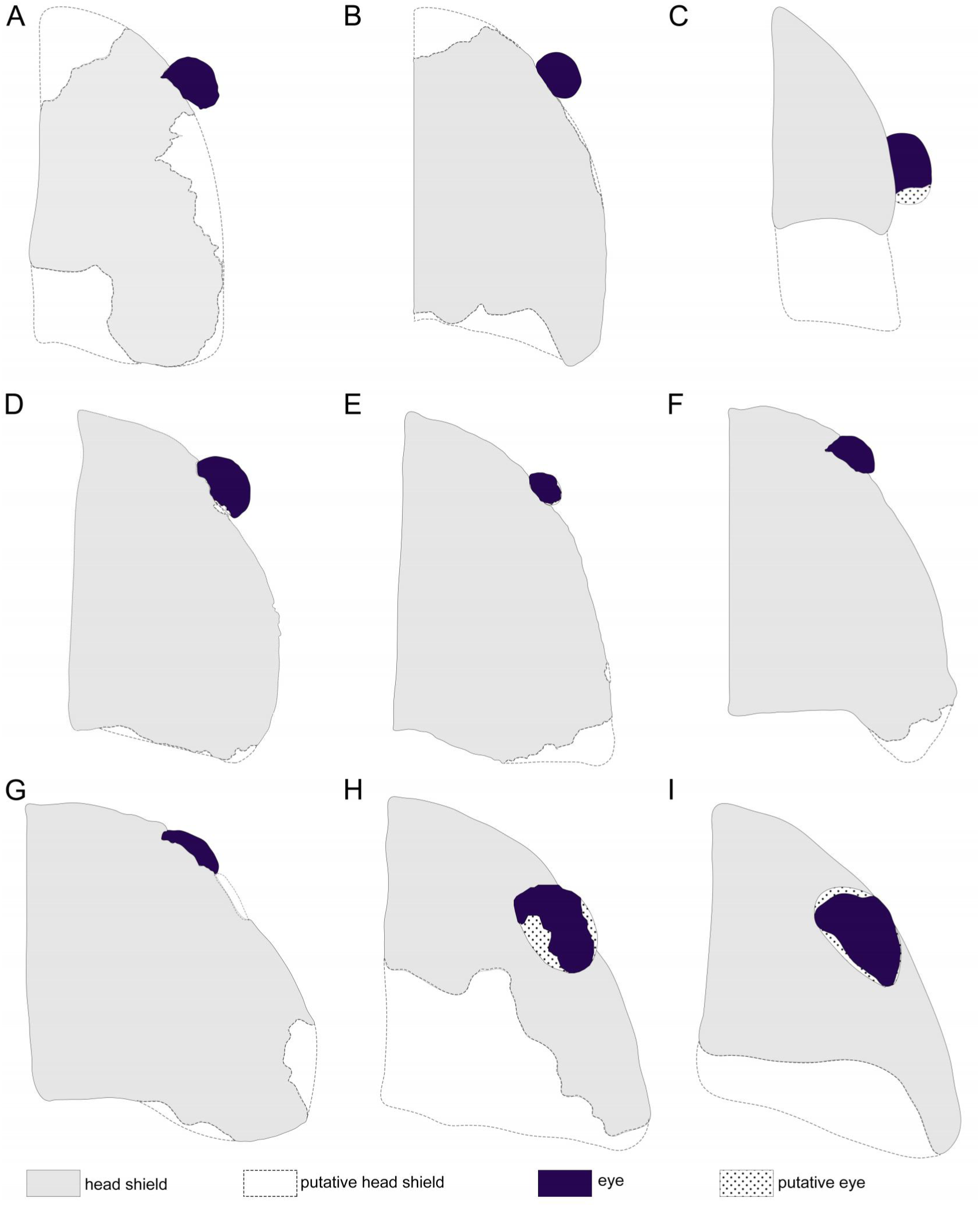
Outline of head shield (half) and eyes in C. eucalla specimens. Gray area indicates head shield, dashed lines indicate the putative head shield, deep purple indicates eye, doted area indicates putative eye. A YKLP 17371. B YKLP 17368. C. YKLP 17366. D YKLP 17362 (flipped horizontal). E YKLP 17367. F YKLP 17376 (flipped horizontal). G YKLP 17365. H YKLP 17373. I YKLP 17372 (flipped horizontal). Not to scale.

**Figure 3—figure supplement 1.**
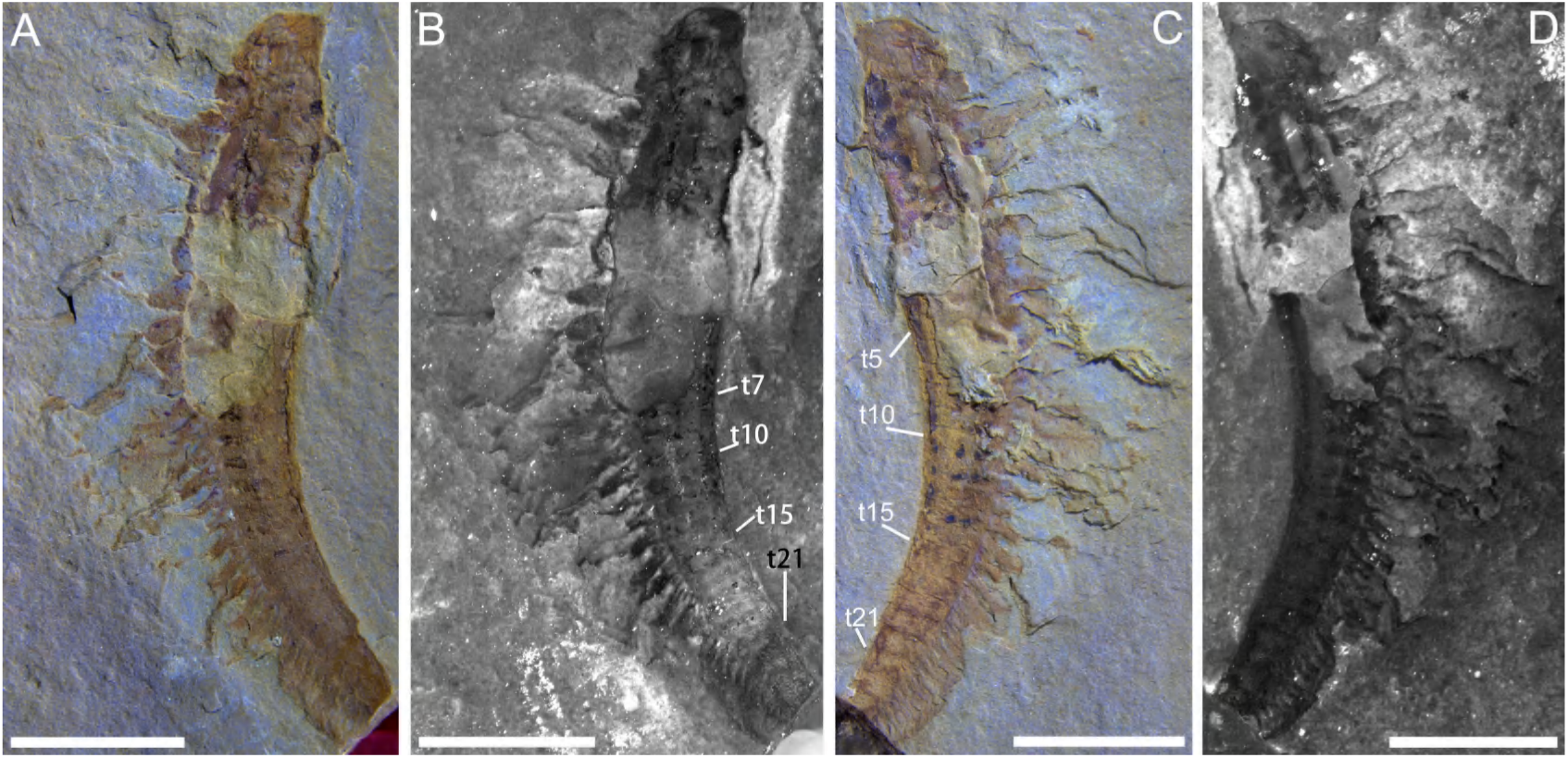
Macro-photographic (A and C) and fluorescent (B and D) images of C. eucalla (YKLP 17369). A and B YKLP 17369a, dorsal view. C and D YKLP 17369b. Abbreviations: t*n*, tergite number. Scalebar: 5 mm.

**Figure 4—figure supplement 1.**
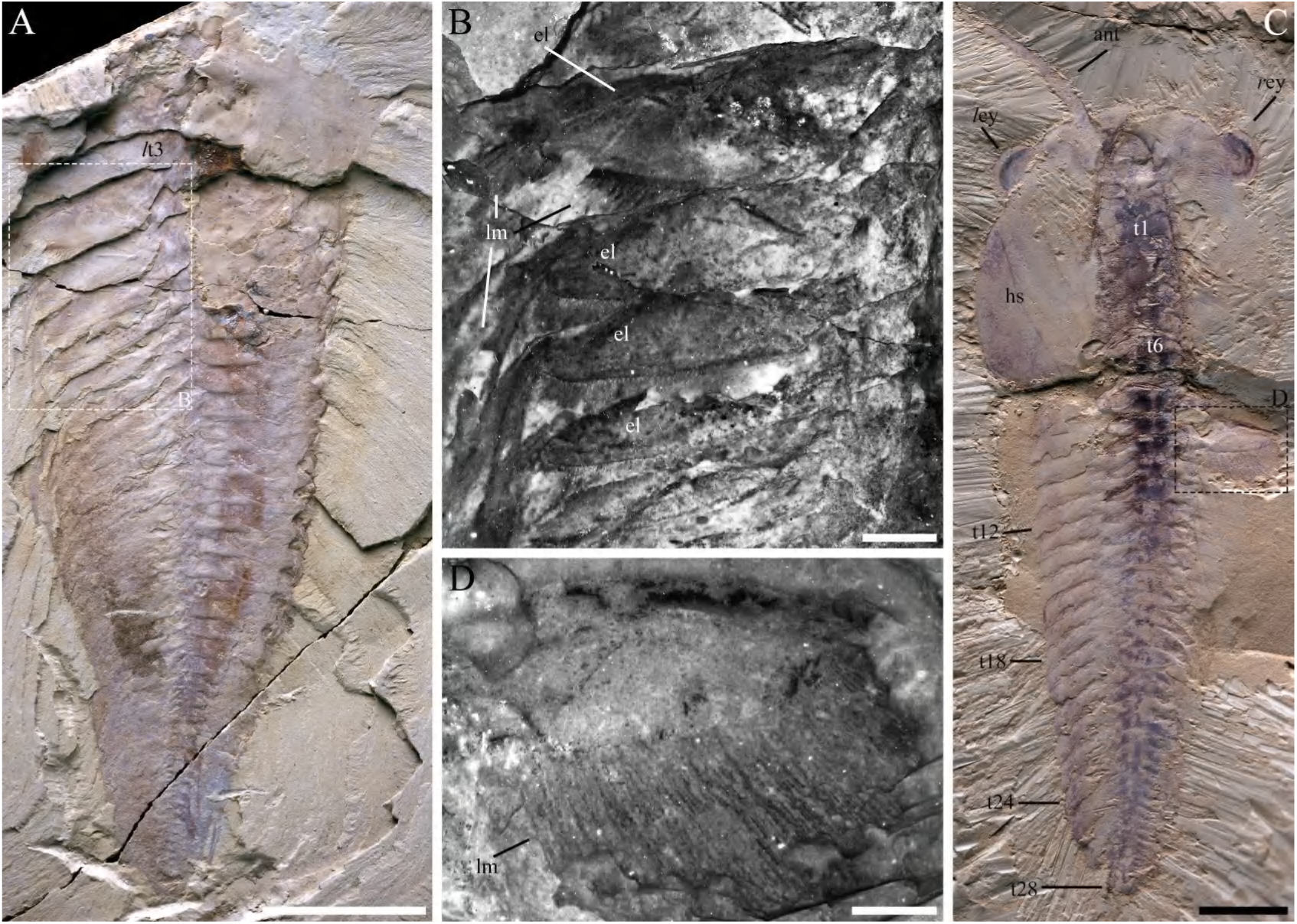
Macro-photographic (A, C) and fluorescent (B, D) images of C. eucalla (YKLP 17369). A YKLP 17374, dorsal view. B close-up of trunk appendages within the dashed box in A. C YKLP 17362, dorsal view. D close-up of a trunk exopod within the dashed box in C. Abbreviations: el, exopod lobe/shaft;lm, lamella(e); hs, head shield; t*n*, tergite number; *l*, left; *r*, right. Scalebar: 1 cm in A, 5 mm in C, 2 mm in B, 1 mm in D.

**Figure 6—figure supplement 1.**
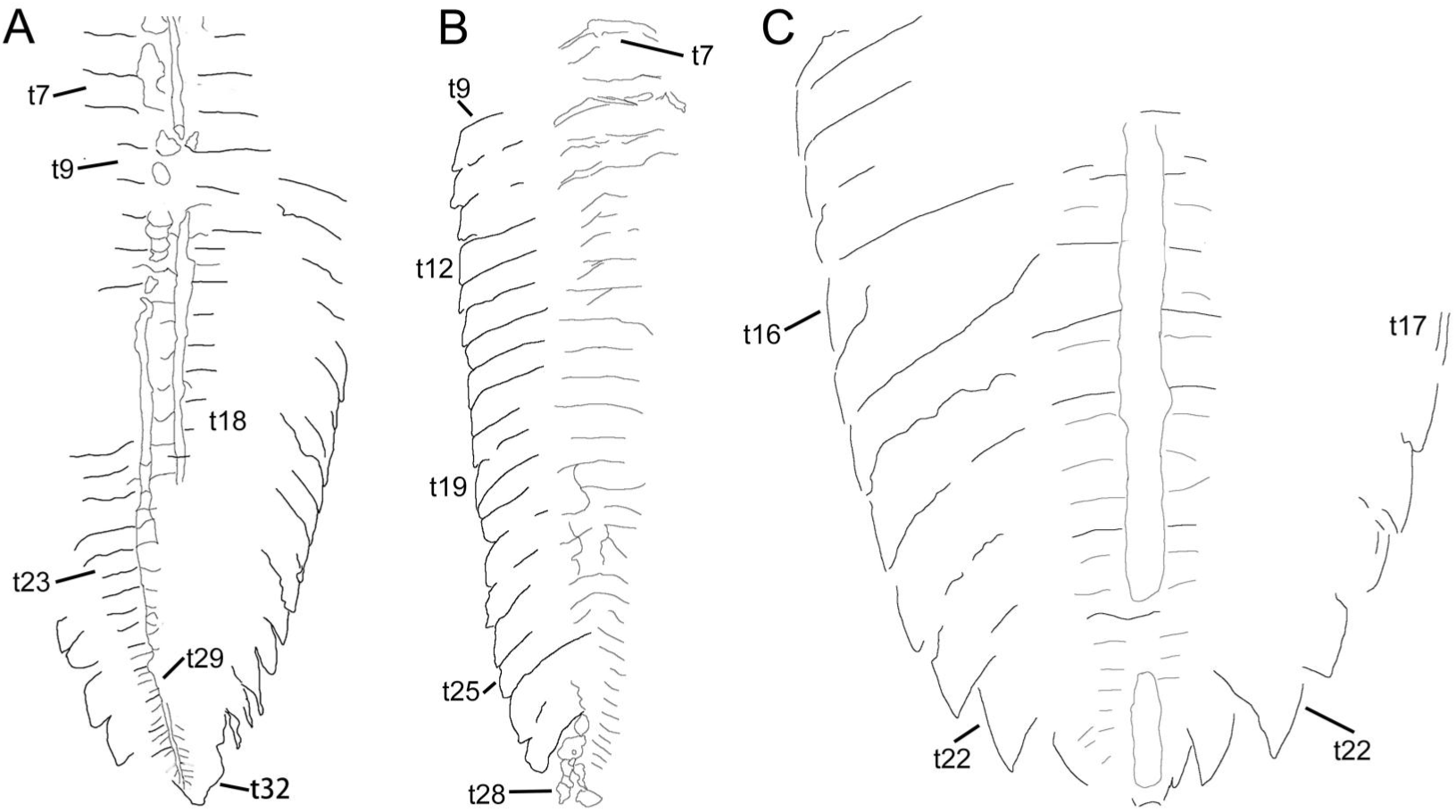
Outline of trunk tergites and ventral segments (gut trace or appendage boundaries). A YKLP 17361. B YKLP 17362. C ELRC 18505. Abbreviations: t*n*, tergite number. Scalebar: not to scale.

**Figure 9—figure supplement 1.**
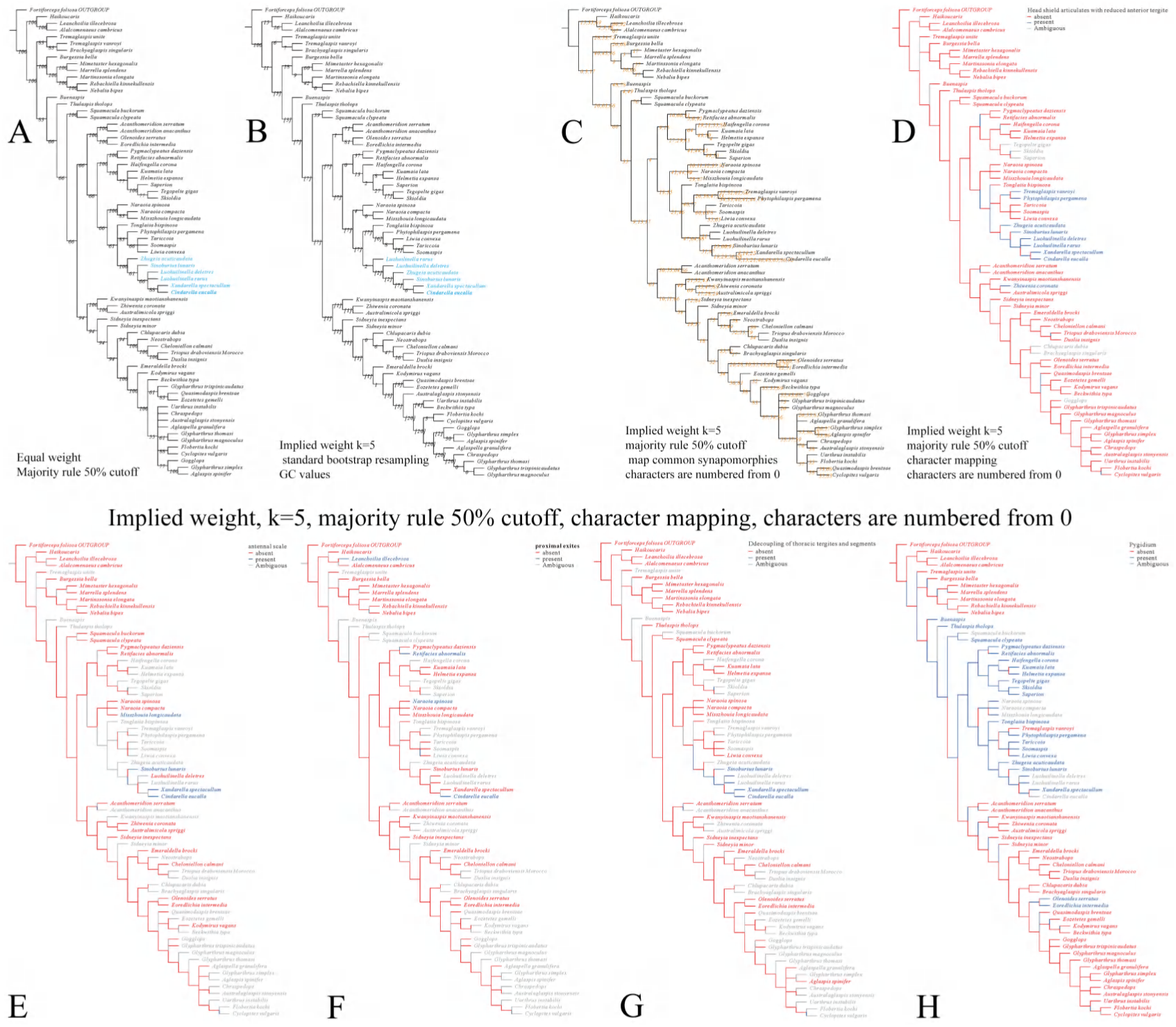
Results of parsimony analysis using TNT v1.6.. A Equal weight parsimony, 50% cut-off majority-rule consensus tree (283 steps, consistency index (CI) = 0.382, retention index (RI) = 0.700, best score = 279) computed from the 18 most parsimonious trees, branch labels indicate clade frequencies (>50%) across the set of most parsimonious trees. B Implied weight parsimony with k=5, nodal support (GC values) was evaluated using standard bootstrap resampling with 2000 replicates. C Implied weight parsimony (k = 5), 50% cut-off majority-rule consensus. Synapomorphies mapped, character numbering starts at 0. D–H Implied weight parsimony (k=5), 50% cut-off majority-rule consensus, character mapping, characters are numbered from 0. D Small tergite(s). E Antennal scale. F Proximal exite. G Decoupling of thoracic tergites and segmenta. H Pygidium.

